# Climate risks to carbon sequestration in US forests

**DOI:** 10.1101/2021.05.11.443688

**Authors:** William R.L. Anderegg, Oriana S. Chegwidden, Grayson Badgley, Anna T. Trugman, Danny Cullenward, John T. Abatzoglou, Jeffrey A. Hicke, Jeremy Freeman, Joseph J. Hamman

## Abstract

Forests are currently a substantial carbon sink globally. Many climate change mitigation strategies rely on forest preservation and expansion, but the effectiveness of these approaches hinges on forests sequestering carbon for centuries despite anthropogenic climate change. Yet climate-driven disturbances pose critical risks to the long-term stability of forest carbon. We quantify the key climate drivers that fuel wildfire, drought, and insects, for the United States over 1984-2018 and project future disturbance risks over the 21st century. We find that current risks are widespread and projected to increase across different emission scenarios by a factor of 4-14 for fire and 1.3-1.8 for drought and insects. Our results provide insights for carbon cycle modeling, conservation, and climate policy, underscoring the escalating climate risks facing forests and the need for emissions reductions to mitigate climate change.

## Main text

Earth’s forests play a fundamental role in the global carbon cycle and currently comprise a substantial carbon sink, sequestering up to 25% of human carbon dioxide emissions annually (1, 2). Governments, corporations, and non-governmental organizations have shown widespread and growing interest in leveraging forests as “nature-based climate solutions” to sequester and store carbon (C) as part of meeting climate policy goals, due to their role as a C sink and additional co-benefits for biodiversity and ecosystem services (3–6). Yet significant scientific gaps remain that greatly limit the effective use of forest-based climate solutions in an evidence-based climate policy framework. Crucially, to be used for climate mitigation, forests must achieve some level of “permanence” whereby they store C for long periods of time (7, 8). Although fossil C emissions persist in the atmosphere for hundreds to thousands of years (9), many public and private policy systems only require forest carbon to last for up to 100 years (8).

The future of forests in a rapidly changing climate is highly uncertain (10, 11). In particular, increasing climate stresses and disturbance could reverse forests’ C sink to a C source and undermine the potential of forests as a climate solution (7, 12–14). For example, an unprecedented and climate-fueled bark beetle outbreak in Canada drove immense swaths of tree mortality and reversed an entire region of boreal forest from a C sink to a C source over a decade with large implications for climate policy (15, 16). In addition to insect outbreaks, wildfires and drought stress have been widely documented as prominent risks because they strongly regulate forest C sinks and are likely to increase in future climates (17–23), which may increasingly compromise regional or continental C sinks. Thus, it is essential to rigorously quantify and understand drivers of historical risks and use this understanding to forecast future climate-driven risks for forest permanence (24–27).

Rigorous forest climate risk assessment is crucial for climate policies and programs relying on forest carbon uptake and storage, yet continental-scale risk assessment is currently lacking and urgently needed (21, 27, 28). Spatial quantification of risks can inform forest protocols in climate policy by ensuring that climate risks are adequately considered in program design — for example, through the construction of “buffer pools” and other insurance mechanisms — and can guide forest project development and conservation (29). However, current forest offset protocols tend to include fixed, spatially invariant risks that do not incorporate future climate impacts and likely underestimate the integrated risks to forests over a 100 year time-horizon (27).

In this paper, we combine forest inventory data across United States (US) forests, remote-sensing data of wildfires, high resolution climate data, and downscaled climate model projections to assess climate-sensitive risks for forests in the US. We first quantify how forest structure and climate anomalies mediate major climate-related risks to US forests from wildfire, drought, and insects. We then model the spatial patterns and magnitudes of these risks over the historical record. Finally, we use downscaled future climate data to project how these risks might evolve in the future due to climate change, revealing where forests are likely to be the most vulnerable in the 21st century.

The fire risk model was based on satellite remotely sensed wildfire burn area from 1984-2018 (Methods). The model reliably predicted historical fires (cross-validated AUC: 0.88), capturing interannual variability (cross-validated *R^2^: 0.62*), seasonal patterns (e.g. spring risk in the southeastern US and fall risk in the western US; cross-validated *R^2^: 0.92*), and spatial patterns (cross-validated *R^2^*: 0.049) (Figure 1A-B, Figure S1). The model captured the spatial patterns of more prevalent fire across the western US, in particular in California and the northern Rocky Mountains, in agreement with previous work (30).

**Figure 1.**
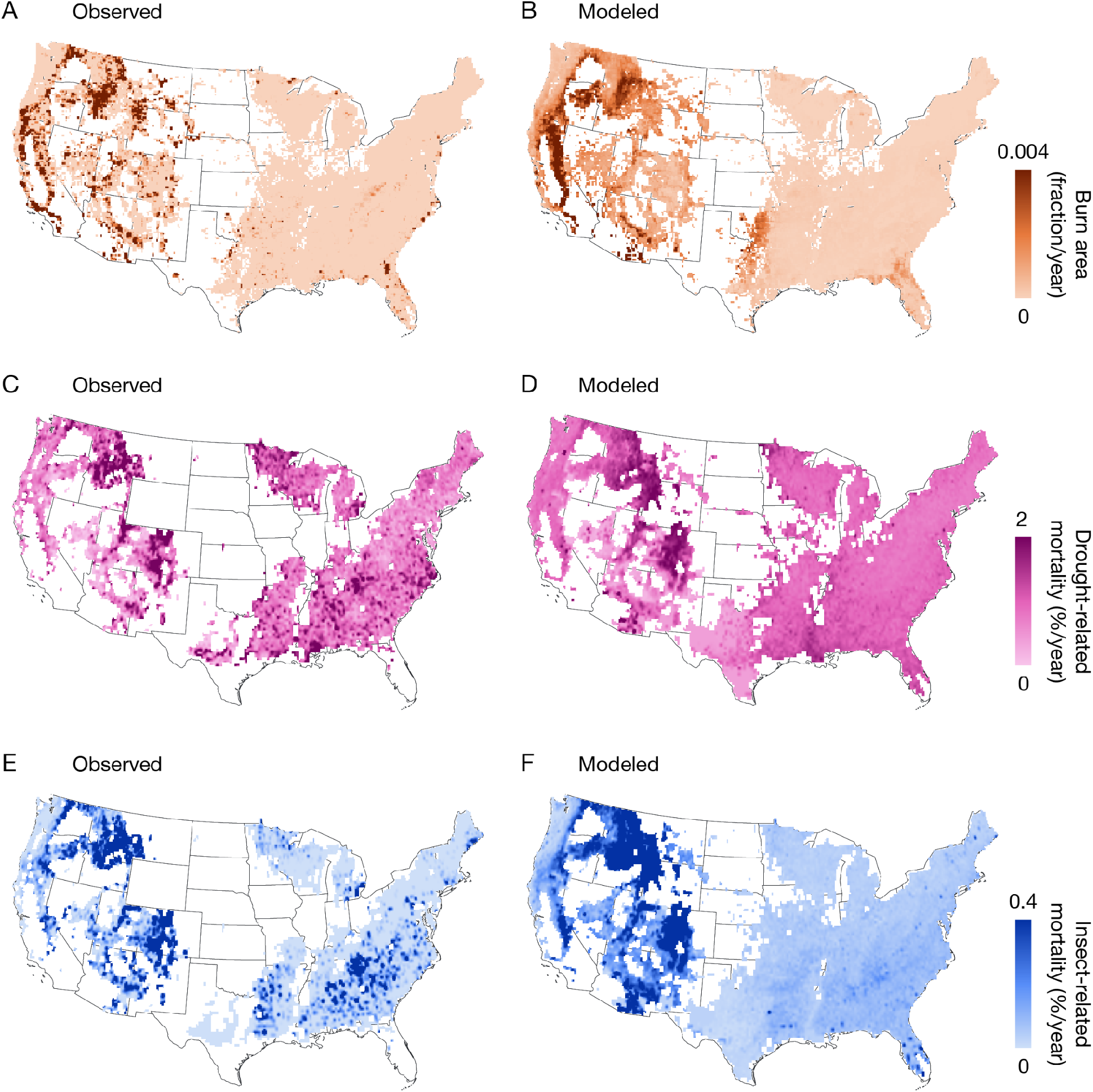
Observed (left) and modeled historical (right) risk maps for fire (A), drought (B), and insects (C) reveal widespread and spatially varying risks. For each impact risk model (B, D, F), anywhere shaded is considered forested. The forest mask for fire differs slightly from those used for insects and drought due to different input data. Data gaps in forest inventory in WY and OK preclude observed risk calculations in drought and insect models (C, E).structure and climate anomalies mediate major climate-related risks to US forests from wildfire, drought, and insects. We then model the spatial patterns and magnitudes of these risks over the historical record. Finally, we use downscaled future climate data to project how these risks might evolve in the future due to climate change, revealing where forests are likely to be the most vulnerable in the 21st century.

The drought- and insect risk models were based on US forest inventory data that tracks tree mortality over >100K long-term plots. Historical patterns of drought- and climate-stress-driven mortality were highest across the western US and intermountain West, which was captured in the mortality model (cross-validated spatial *R^2^: 0.18*; Figure 1C-D) and is consistent with recent studies and other independent metrics (31), Figure S2). The inclusion of forest physiological metrics for drought tolerance, specifically community-weighted plant hydraulic traits (32), substantially improved the predictive accuracy of the drought mortality models (ΔAIC<<-10), agreeing with drought-physiology studies (33). Observed historical permanence risks to US forests from insect-driven mortality were highest in the Rocky Mountains and modeled risks captured the key broad spatial patterns in risks (cross-validated spatial *R^2^: 0.31;* Figure 1E-F). These observed and modeled insect risks showed strong spatial agreement with an independent continent-wide insect outbreak dataset (Figure S3) and reflect scientific understanding from previous western US studies (17, 19, 34). Further, the climate sensitivities of insect mortality for several western US pine species with the highest historical insect-driven mortality were consistent with estimates in the literature (Fig. S4) (17, 35). In sum, these results provide an unprecedented synthesis of fire-, drought-, and insect-driven climate risks to forest permanence in an open source dataset available at continental scales.

We combined the historical risk models with a new downscaled 4-km CMIP6 climate dataset (see Methods for dataset description) to project fire, drought, and insect risks over the 21st century for three shared socioeconomic pathway (SSP) scenarios (Fig. 2). Under all future climate scenarios, fire risks are projected to increase substantially throughout the 21st century (Figure 2A). Future risks increase similarly across scenarios through mid-century, but diverge by 2050. By 2080-2099, in a world with high fossil fuel development (SSP5-8.5), US-averaged fire risk is projected to increase 14-fold compared to historical CMIP (average 1990-2019) values. In lower emissions scenarios increases are still dramatic: the multi-model mean projects either 4-fold (SSP2-4.5) or 9-fold (SSP3-7.0) increases. Projected drought risks increased substantially and varied by emissions scenario with average mortality increases by a factor of 1.8 in SSP5-8.5, 1.5 in SSP3-7.0, and 1.3 in SSP2-4.5 by 2080-2099 (Figure 2B). Future insect risk projections indicated a 1.7-fold increase US-wide in SSP5-8.5, 1.4-fold in SSP3-7.0, and 1.2-fold in SSP2-4.5 by 2080-2099 (Figure 2C). All three climate-sensitive risks showed large differences across climate models, although the relative ranking of risk by SSP was consistent by the end of the century. The substantial differences between high and low emissions SSPs emphasizes the critical importance of climate policy to mitigate climate risks to US forests.

**Figure 2.**
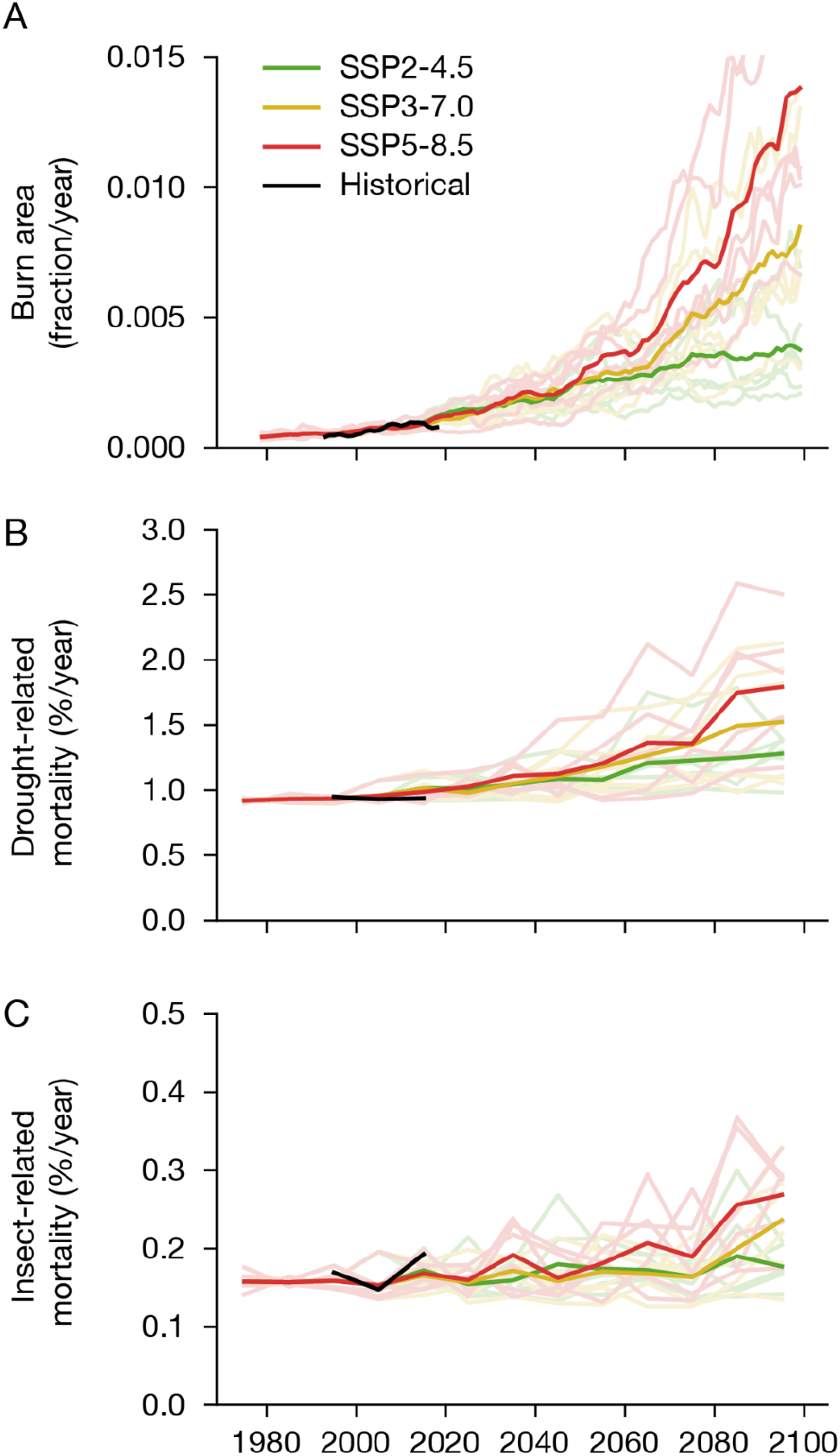
Projected 21^st^ century risks for fire (A), drought (B), and insects (C) averaged across the US. Simulations from each GCM shown as transparent lines, colored according to the three shared socioeconomic pathway (SSP) climate scenarios. The multi-model mean for each SSP is shown opaque. Historical modeled simulations are shown in black. Fire risks are calculated with a 10-year centered moving average, while drought-related and insect-related are presented as decadal averages.

We then conducted a risk assessment to quantify which regions and forests are likely to experience the highest climate-sensitive risks in the 21st century (Figure 3, representing the average of 2080-2099). By the end of the 21st century, high levels of risk which used to be confined to pockets in California and the intermountain western US are projected to expand across the entire western US (Figure 4). While these risks are substantially mitigated by emissions reductions (SSP2-4.5, Figure 3A), risks are still projected to increase dramatically in regions like the Great Plains in the central US and southeastern US (Figure 4). Patterns of increased risks are spatially consistent with previous work that found stronger relative changes in regions of comparatively lower historical risk (20).

**Figure 3.**
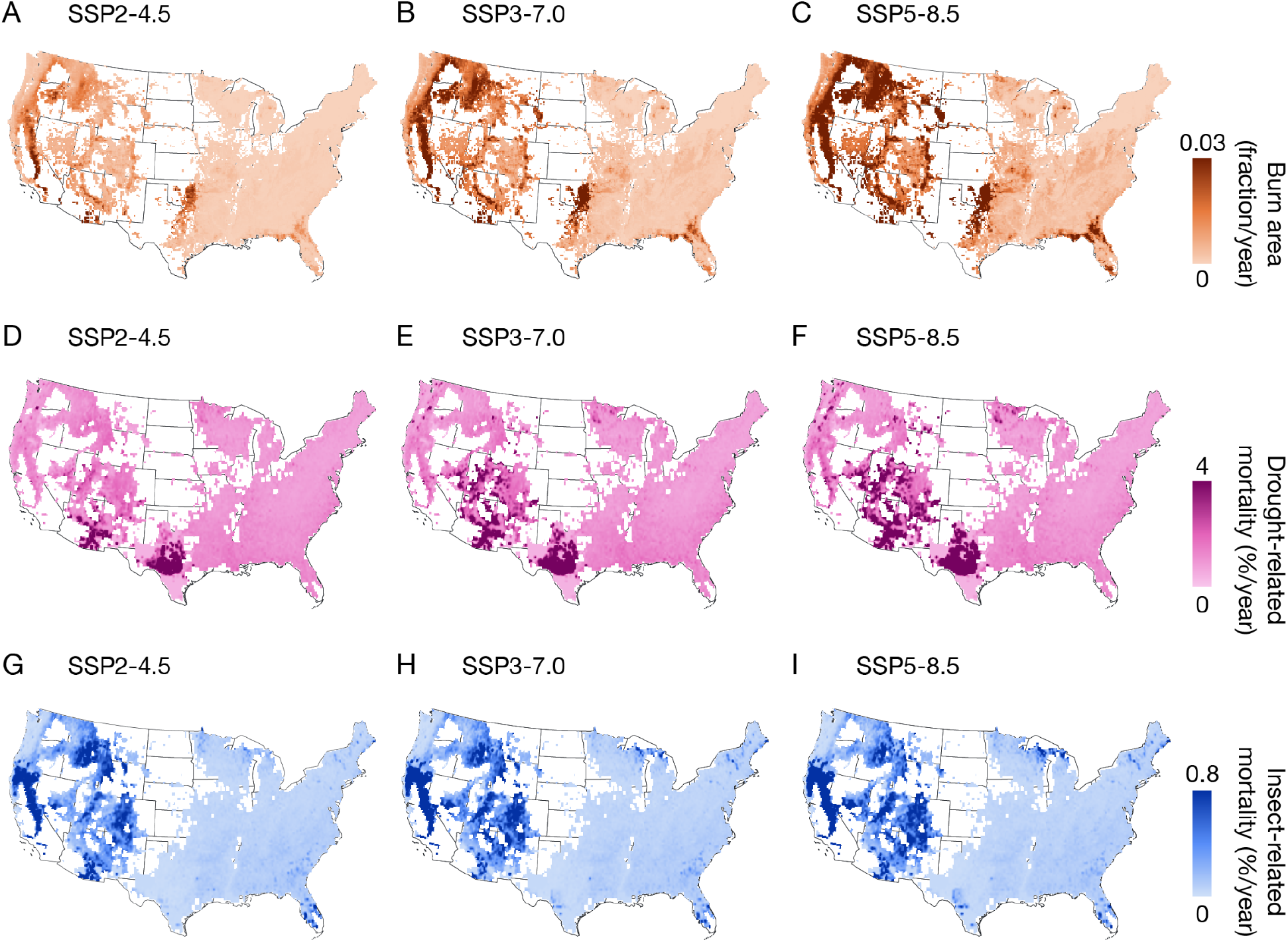
Risks for fire, drought, and insects (rows) averaged over the 2080-2099 period, separated by shared socioeconomic pathway (SSP) climate scenario (columns). Note that colorbars are substantially expanded relative to those in Figure 1 in order to visualize future projections that exceed historical risks.

**Figure 4.**
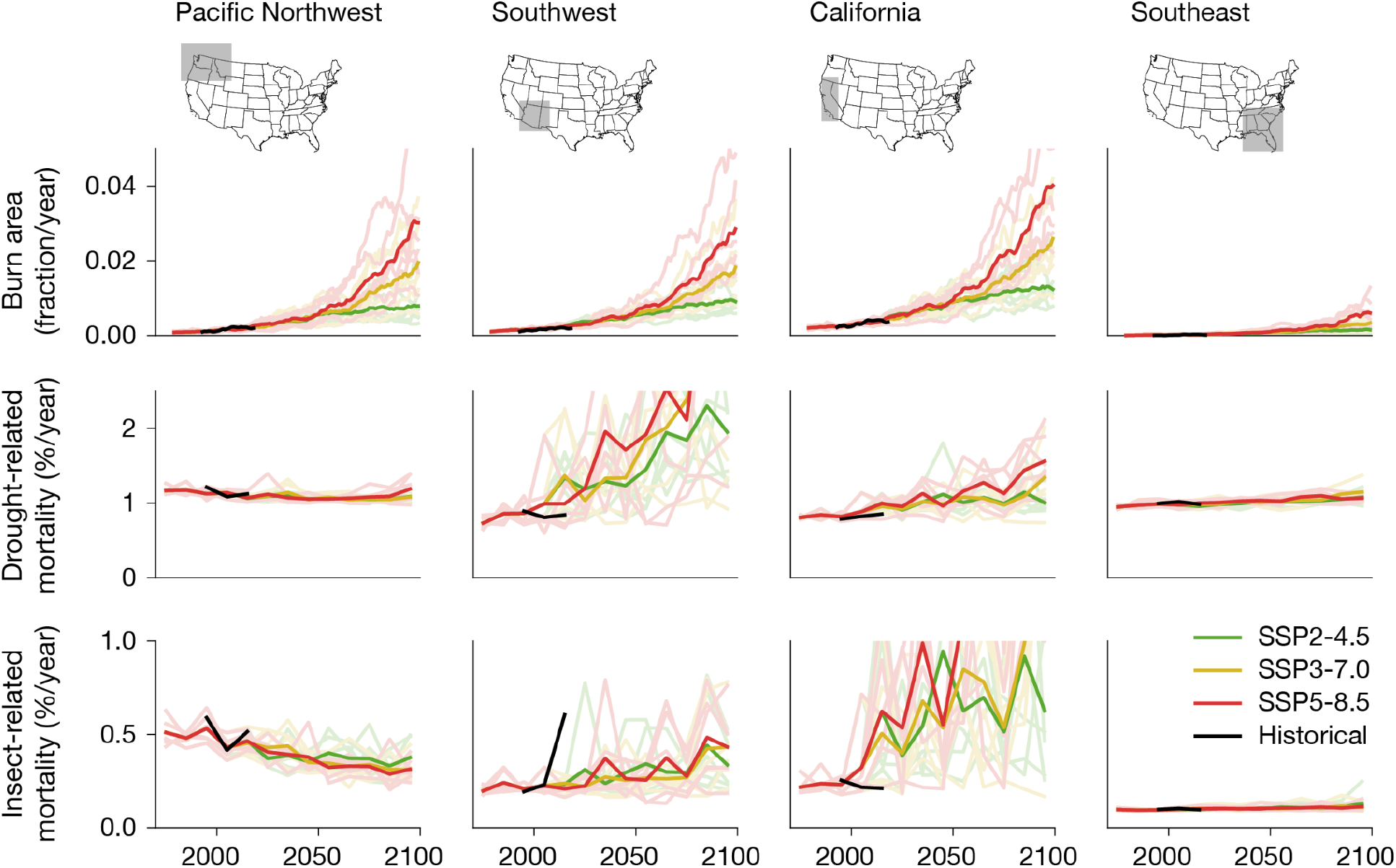
Regionally-averaged time series of forest risks from three impacts: fire (top), drought (middle), and insects (bottom). Regions of interest shown as gray boxes in the maps at the top of each column. As in Figure 2, simulations from each GCM simulation are shown as transparent lines, colored according to the three shared socioeconomic pathway (SSP) climate scenarios. The multi-model mean for each SSP is shown opaque. Historical region-averaged modeled simulations are shown in black. Fire risks are calculated with a 10-year centered moving average, while drought-related and insect-related are presented as decadal averages.

Future drought risks increased most across broad swaths of the intermountain and southwestern US, California, and western Texas, although parts of the eastern US and the upper midwestern US also exhibited increased drought mortality risk (Figure 3B; Figure 4). Projected insect risk to forest permanence was highest across the Rocky Mountains in the intermountain western US, Sierra Nevada mountains in California, and parts of the northern Midwest (Figure 3C; Figure 4). The spatial patterns of these projections are consistent with previous projections for a few major insect species (36) and overall magnitude is similar to coarse-level ecoregion projections in parts of the western US (37) (Figure 4). We note that drought and insect model projections were only made for forest types where models showed skillful cross-validated performance (i.e. AUC>0.6) and thus lower risk in some regions (e.g. southern pine beetle risk in the southeastern US (38)) may reflect data and model limitations rather than inherently lower risks.

These climate-sensitive risk maps and projections provide spatial quantification and uncertainty assessment across climate models, climate scenarios, and risk models (Figure S5) that enables rigorous risk management and conservation decisions.

To support these aims, all data and code underlying these models is publicly available (https://doi.org/10.5281/zenodo.4741329, https://doi.org/10.5281/zenodo.47413334), and can be easily accessed and visualized via a web portal (https://carbonplan.org/research/forest-risks). We note that, as with any analysis, these projections are subject to several uncertainties and caveats. In addition to uncertainties in underlying Earth system models and statistical climate downscaling approaches (Methods), these projections use empirical models based on static forest composition and structure over the 1984-2018 period. Thus, these projections do not account for shifts in forest composition or distribution, interactions among risks, and only partially include CO_2_ effects on plant drought stress (Methods). In particular, large-scale impacts of fires, drought, or insects could substantially reduce biomass, and thus risk, as a negative feedback, although this is not thought to be likely before the 2050s (20, 21, 25). The risk projections also do not include impacts of land-use management, considered to be a strong potential lever in fire risk (39) and to a lesser degree drought and insect risks. We note, however, that there are also many reasons that these risk projections may be conservative or underestimates, including projections made only for strong historical models, frequent non-linear impacts of drought and insects that may not be well-characterized in inventory data, and novel pests and pathogens (Methods).

Climate-sensitive risks to US forests have major impacts on forest C cycling and climate change feedbacks, which is why it is critical to quantify forest permanence risks for conservation and climate policy efforts. Tree mortality and disturbance are among the largest uncertainties in current land surface and vegetation models (40–42) and better large-scale historical datasets are needed for these models. Thus, the disturbance risk and mortality maps and their climate sensitivities derived here provide a foundation for advancing C cycle models. Our results reveal that US forests are very likely to experience increasing risks from climate change that could undermine their C sequestration potential, an important factor that should be considered in climate change mitigation policy. First-order estimates of the policy-relevant US-wide 100-year integrated risk of “reversal” (i.e. loss of carbon) were 8% for fire, 19% for drought, 20% for insects based on historical data alone and were 40% for fire, 26% for drought, and 27% for insects in SSP5-8.5, compared to the 2-4% stationary reversal risk widely used for each disturbance in many forest offset protocols. Thus, our results make clear that 21st century reversal risks are massively underestimated in current policies and must be updated to achieve climate mitigation goals. Taken in sum, our results strongly underscore the needs for reducing greenhouse gas emissions to mitigate climate change given the increasing risks of climate change undermining forest permanence and natural climate solutions.

## Materials and Methods

### Datasets

Several datasets were used for fitting all our models. Here we describe each dataset, and how we preprocessed it for the purpose of analysis.

#### MTBS

The Monitoring Trends in Burn Severity (MTBS) dataset includes 30-m annual rasters of burn severity as well as burn area boundary polygons for individual fires (43). The dataset covers fires from 1984-2018 and includes fires larger than 202 ha (404 ha in the Western U.S.) for the continental U.S., Alaska, Hawaii, and Puerto Rico.

For our analysis, we first created monthly rasters of burn area on a 30-m grid by rasterizing the polygons of all fires from the MTBS database within individual months. We then additionally masked each monthly raster so as to only include pixels corresponding to moderate or high burn severity. (That assignment of burn severity was based on the annual thematic raster from the corresponding year). These monthly rasters were then resampled to a 4-km Albers equal area grid using averaging, in accordance with the spatial resolution of our meteorological data (described below). Any pixels identified as non-data (due to LANDSAT artifacts in the raw MTBS record) were excluded during resampling. This preprocessing results in a raster where each pixel encodes the fraction of that 4-km grid cell that burned during the corresponding month.

Notebooks and scripts to reproduce the MTBS rasterization process can be found at https://github.com/carbonplan/data/tree/main/scripts/mtbs. Individual 30-m monthly rasters are available at https://carbonplan.blob.core.windows.net/carbonplan-data/raw/mtbs/conus/30m/area/{YYYY}. {MM}.tif in Cloud Optimized GeoTIFF (COG) format and a 4-km monthly spatio-temporal raster is available at https://carbonplan.blob.core.windows.net/carbonplan-data/processed/mtbs/conus/4000m/monthly.zarr in Zarr format.

#### FIA

The Forest Inventory and Analysis (FIA) dataset is a nationwide survey made up of an extensive series of long-term forest monitoring plots that the USDA Forest Service uses to monitor the growth, mortality, and overall health of US forests. Raw FIA data were downloaded from the FIA Data Mart in CSV format on August 6th, 2020.

Our analysis relied on two components of the FIA database: tree level measurements and condition-level measurements. A condition is a homogenous subdivision of an FIA plot of the same land cover type (e.g. forest vs non-forest). From the raw, per-tree measurements, we calculated a series of condition-level summary statistics, including total living and dead basal area, and combined these with other condition-level summaries from the database. We aligned conditions across repeated measurements of the same plot. For all analyses, we only included conditions with a “condition proportion” greater than 0.3, which means that the condition represented at least 30% of its corresponding plot, following a recent similar analysis (32). We tested to ensure that historical drought and insect model results were robust to this threshold by varying this threshold between 0-30% and found very similar patterns (e.g. *R^2^*>0.95 for all threshold modeled mortality comparisons).

Preprocessed versions of these data for individual states in Parquet format are available at https://carbonplan.blob.core.windows.net/carbonplan-data/processed/fia-states/long/{ST}.par quet where {ST} is a state abbreviation in lowercase.

#### NFTD

The National Forest Type Dataset, based on FIA data, is a collection of rasters which encode the most common forest type and forest group at 250 m (44). For each forest group, we constructed a raster resampled to the same 4-km grid encoding, for each grid cell, the fraction of that forest group present.

#### NLCD

The National Land Cover Database (NLCD; (45)) includes 30-m land cover classification rasters at 2-3-year intervals (from 2001-2016). Separate raster datasets are available for the contiguous U.S. and Alaska.

Our processing of the NLCD dataset included downsampling the categorical data from 30-m to 4-km resolution. For this study we considered regions within 20 land cover categories of interest (11, 12, 21, 22, 23, 24, 31, 41, 42, 43, 51, 52, 71, 72, 73, 74, 81, 82, 90, 95), representing forested, shrubland, herbaceous, and cultivated areas. We extract a binary coverage mask for each type. The downsampling process thus results in fractional coverage 4-km rasters for each of the 20 land cover categories.

See the NLCD downsampling and reprojection notebook for specific details of the process (https://github.com/carbonplan/data/tree/main/scripts/nlcd).

#### TerraClimate

The TerraClimate dataset provides monthly climate and climatic water balance terms (46). The dataset extent includes all land areas on a global 1/24th degree grid from 1958-2019. To process the TerraClimate dataset, we regridded the data from its global 1/24th degree grid to our 4-km project grid using xESMF’s bilinear interpolation.

#### CMIP6

The Coupled Model Intercomparison Project Phase 6 (CMIP6; (47)) was used as the basis for our future climate projections, though we also evaluated the model on historical CMIP forcings to verify the model’s ability to transfer from TerraClimate to CMIP forcings. We extracted monthly mean surface climate data for precipitation, minimum daily temperature, maximum daily temperature, downwelling shortwave radiation, relative humidity, and surface air pressure from the CMIP (historical) and ScenarioMIP (SSP2-4.5, SSP3-7.0, SSP5-8.5) scenarios. For this study, we selected all models that had complete data for one ensemble member in each of the four scenarios. We note that these are merely a sampling of a growing body of future climate simulations and expanding the ensemble size as more simulations become available will improve the understanding of uncertainty around our projections.

| Model | Member |
| --- | --- |
| ACCESS-ESM1-5 | r10i1p1f1 |
| MRI-ESM2-0 | r1i1p1f1 |
| MPI-ESM1-2-LR | r10i1p1f1 |
| CanESM5-CanOE | r3i1p2f1 |
| ACCESS-CM2 | r1i1p1f1 |
| MIROC-ES2L | r1i1p1f2 |

### Downscaling and deriving hydrologic variables

The CMIP6 data described above were further processed to support our climate impacts analysis. The model output was first regridded to our 4-km project grid using xESMF’s bilinear interpolation. Then the data were bias-corrected and run through a simple hydrology model, both steps described in detail below. For all climate data processing, we used the years 1970-2000 (inclusive) as our climate normal period.

#### Quantile Mapping

Quantile mapping is widely-used in the climate downscaling literature with a variety of implementations depending on the application (48–51). For this study, we performed a simple form of single-variable trend-preserving quantile mapping (50, 52) to correct for persistent biases in the raw CMIP6 climate model output, matching the statistical distribution of the TerraClimate dataset on a gridcell by gridcell basis. Using the Scikit-downscale package (Hamman and Kent, 2020), the quantile mapping procedure was implemented for each variable and grid cell as follows. The process begins by extracting the three timeseries:

*X: historical (i.e. CMIP)*
*y: observed (i.e. TerraClimate)*
*X’: modeled (i.e. CMIP or ScenarioMIP)*

and by removing their trends. Empirical CDFs are then calculated for all three timeseries using Cunnane plotting positions. The CDF from *X*’ is then interpolated onto the CDF of *X* using one-dimensional piecewise linear interpolation, thus correcting for statistical differences between the *X* and *X*’. The mapping from *X* to *y* is performed by again using one-dimensional piecewise linear interpolation. Finally, the trend from *X*’ is reimposed to the resulting vector of quantile mapped data.

Though this method is performed on monthly data, we still take care to handle extreme values in a reasonable way. As implemented in our study, we take the common approach of applying the proportional differences between *X*’ and *X* for all values outside the bounds of the training data.

#### Hydrology

We used multiple hydroclimatic variables from TerraClimate in the development of the individual risk models (described below). Here we reimplemented the core components of the simple hydrologic model from (46), the code for which is available here [https://github.com/carbonplan/cmip6-downscaling/blob/main/cmip6_downscaling/disagg/terraclimate.py]. We used the PDSI implementation of the climate_indices software package (53). Surface elevation and available water content data was taken directly from the TerraClimate dataset. We note that the PET-related variables contain a “CO_2_ adjustment” factor that aims to account for the effects of CO_2_ on decreasing water loss through declines in surface (e.g. stomatal) conductance, which enables a more rigorous assessment of drought stress projections in future climates.

### Risk Models

#### Fire

We developed a statistical model predicting burn area as a function of climatic variables. Our work is inspired by, and builds, on previous fire risk estimation efforts (30). Many of the methods are similar, though updated with more recent data (through 2018 rather than 2010).

The model was fit to historical fire data from the Monitoring Trends in Burn Severity (MTBS) database (described above). Our dependent variable was thus a (420) x (1209) x (783) raster of monthly burn area per month, many entries of which were 0. As regressors, we considered both temporally-varying climatic variables as well as time-invariant variables to account for heterogeneity in vegetation. Our primary climatic variables were mean temperature and summed precipitation derived from the TerraClimate dataset (described above) for the historical period matched to the MTBS record as well as climatic water deficit (CWD). This resulted in a (3) x (420) x (1209) x (783) regressor raster.

For vegetation, we used the National Forest Type Dataset (described above). To limit the number of variables and prevent overfitting, spatially sparse forest groups were aggregated into supersets by combining any forest group with spatial area less than 1.76M ha with the most spatially similar, larger forest group (areas ranged from 51k ha to 67M). This grouping removed 8 out of 25 groups, and had little effect on model behavior. Thus, the result of our preprocessing was a (17) x (1209) x (783) forest group regressor raster.

We fit a “hurdle” regression model to predict burn area as a function of climate and vegetation variables. This model jointly predicts the probability of a non-zero value and, if a non-zero value is present, its continuous value (54). Intuitively, this model can be thought of as combining a classifier (“was there fire?”) and a regression (“if there was fire, how large was the burn area?”). For computational reasons, all datasets were coarsened by a factor of 4x prior to fitting.

We formally represented the hurdle model using a sequence of two generalized linear models: a Binomial model with logit link function predicting zero vs non-zero values, and a linear Gaussian model with normal link function predicting burn area in the locations where it was non-zero. We implemented the hurdle model in Python using scikit-learn by combining the LogisticRegression and LinearRegression methods (55).

In addition to the variables described above, we included two additional regressors to better capture inter-annual trends, which the model might otherwise ignore given that variation within years is substantially greater than variation across years. To create this regressor we calculated a US-average temperature and precipitation time series and then calculated, on a monthly rolling basis, the 12-month maximum temperature and precipitation timeseries. We used the resulting timeseries as a regressor which was the same at all locations but varied over time. Conceptually, the temperature regressor provides a measure of longer term drought stress than the short term monthly timeseries. The precipitation regressor captures the sharpness of precipitation regimes, which has been shown to amplify wildfire risk in California (56). In practice, including these extra regressors improved overall model performance only slightly, but allowed the model to better reproduce both monthly trends, interannual variability, and the observed increase in burn area over the observation period, a large fraction of which has been attributed to climate change (see Figures S7–8).

We assessed model accuracy using both area-under-the-ROC-curve (AUC) from the output of the logistic regression portion of the hurdle model, as well as *R^2^* from the linear regression portion. Given the sparse and largely binary nature of the data, especially compared to the insect and drought models, we focus on AUC as the primary performance metric of interest. We report these AUC values obtained using split-halves cross-validation, where the held out set was constructed by sampling years independently, i.e. by holding out a contiguous period of time (Figure S6). We also assessed performance through the model’s ability to reproduce three trends: annual, seasonal, and spatial. Each of these trends corresponds to a “slice” or “average” of the spatio-temporal raster — specifically, averaging across months and space (for the annual trend), averaging across years and space (for the seasonal trend), and averaging across time (for the spatial trend). In each case, we computed an *R^2^* between the value computed directly from the data and the model’s prediction allowing for a constant offset difference (Figure S6). While these are not the metrics on which the model is trained, they provide an indication of how well the model captures important and observable patterns in the data. In general, these metrics increased alongside increases in overall model performance.

As an additional form of cross-validation, we considered the ability of the model to extrapolate beyond the period of which it was fit. For each of several years Y, we fit the model on years between 1984 and Y, and predicted on years between Y + 1 and 2018, and computed an *R^2^* as above. Note that, because our calculation of *R^2^* allows for an arbitrary offset, it can be high even when the absolute magnitude of the prediction is off (i.e. when there is a bias), which is especially likely when extrapolating model fits. For that reason, when computing these statistics, we separately compute and report the bias as the fractional difference between the average risk between prediction and observation (Figure S1).

For visualization purposes, predicted monthly burn areas from the model (fraction / month) were summed across months to estimate predicted burn area for each year (fraction / year). When making future projections, we interpret the modeled annual burn area for each grid cell as a uniform annual probability of fire anywhere within that grid cell, i.e. as a fire risk.

#### Drought and Insects

We aggregated FIA data on live and dead basal area from a tree-level to a ‘condition’ level, grouping together conditions representing repeated inventories of the same location. To construct drought and insect risk models, we screened for plots that had at least 2 or more inventory measurements, which enables the estimation of a true mortality rate. We next screened out plots that had a “fire” or “human” disturbance code or a “cutting” treatment code to remove major confounding disturbances.

We estimated the fraction of mortality based on the concept of a census interval, which we define as a pair of measurements in two successive years (t_0_, t_1_). The fraction of mortality is defined as the ratio of dead basal area in t_1_ to the total live basal area in t_0_, which was then normalized by the census length to give annual mortality rates. We computed this ratio separately for each condition. Given that many FIA plots only had one repeated measure (only one census interval), we used the first census interval for all conditions. We modeled ‘drought-driven’ mortality as the mortality that occurred during this census interval (with other confounding mortality drivers excluded, see above) and ‘insect-driven’ mortality using the “agent code” (AGENTCD) tree-level data, where codes of 10-19 indicate insects as the primary causal agent of death. We note that the drought mortality models include mortality from insects, which was a deliberate decision because insects and drought often co-occur and interact to kill trees in many forests across the US and thus cannot be clearly separated (57), although we performed a sensitivity analysis of the drought model when excluding mortality of trees with insect agent codes and observed very similar modeled mortality patterns (*R^2^*=0.60, p<0.0001). Drought and other climate-driven mortality does not have a clear or widely used agent code in the FIA database; instead, drought-driven mortality is often attributed to a wide array of more proximate agents (including insects, disease, weather, and other/unknown); see ref (57) for a detailed discussion. Thus our attribution is that this mortality is likely driven by “climate stress” broadly defined, as we have aimed to remove other major drivers of mortality, notably fire, human disturbance, and account for stand and self-thinning dynamics in model construction (see below). This mortality attribution approach has uncertainties but is generally reasonable and is the standard approach that has been widely used in numerous climate-related FIA studies (32, 57–61).

We fit a statistical model predicting mortality as a function of climatic and stand variables. We formally represented the hurdle model using a sequence of two generalized linear models: a Binomial model with logit link function predicting zero vs non-zero values, and a linear model with beta-distributed link function, which is used for modeling proportions where values are between zero and one, for predicting mortality in the conditions where it was non-zero. The beta-distributed link function for the linear regression was chosen based on inspecting the behavior of the raw data distributions. We implemented the hurdle model in R using the ‘glm’ in the default stats package and the ‘betareg’ package (62).

For each condition, we extracted the mean, minimum, and maximum over the census interval for six annual climate variables that were selected based on their importance in the drought and insect mortality literature: precipitation, temperature, palmer drought severity index (PDSI), potential evapotranspiration (PET), climatic water deficit (CWD), and vapor pressure deficit (VPD) (17, 18, 35, 63, 64). We also extracted the stand age for each condition from FIA and the community-weighted mean and range of the functional trait of the water potential at 50% loss of hydraulic conductivity (P50) from 0.25 degree maps published in a recent study (32). This trait has been widely linked to drought-driven mortality risk in site-level (65, 66) studies and meta-analyses (33). We also included in the mortality models two stand variables of age or age-squared to account for background ecological dynamics such as self-thinning and background mortality, following ref (59). All predictor variables were z-scored across the full dataset for that variable to ensure that variable ranges did not drive model outputs.

Drought and insect mortality models were fit independently to each FIA “forest type,” which was chosen as an intermediate compromise of capturing the diversity of responses across US forests but aggregating above a species-level to enable adequate estimation of mortality levels. To ensure that each forest type had 50 or more condition measurements, we aggregated some sparse forest types into more common ones (59 were so aggregated out of the initial set of 171), leading to 112 initial forest types in our dataset. Because FIA plots are inherently small and may not robustly estimate mortality rates at a plot-level, we subsequently aggregated condition-level mortality rates, age, climate data, and functional traits to a 0.25 degree grid for each forest type. This grid size was chosen through sensitivity analyses to determine the optimal aggregation where the coefficient of variation of mortality rate stabilized but large-scale climate variation was preserved. All drought and insect mortality models were fit on this 0.25 degree grid.

We considered collinearity among predictor variables by examining variance inflation factors. We found that variance inflation factors were too high for comparing mean/min/max of the same variable (e.g. mean vs min vs max annual temperature), but were within acceptable levels (<5) across the six predictor climate variables, stand age, and P50 hydraulic trait. Thus, we conducted a stratified model selection analysis where we fit all possible model combinations with one of each predictor variable (i.e. varying all possible combinations of mean vs min vs max of each climate variable, age vs age-squared, P50 mean vs P50 range) and selected the most parsimonious model via Akaike Information Criterion (67). For all analyses in this paper, we fit the same predictor variables across all forest types to reduce complexity. Thus, individual forest types were not allowed to have separate predictor variables. Model selection analyses were done separately on drought and insect mortality dependent variables.

We examined optimal model complexity by comparing the AIC and *R^2^* of nested sets of models. We compared drought and insect mortality models as a function of: i) a null model of forest type-only (i.e. each forest type would receive only its mean mortality), ii) a null model of mortality as a function of forest type and age only (i.e. no climate predictors), iii) mortality as a function of forest type, age, and climate predictors, and iv) mortality as a function of forest type, age, climate, and functional traits. We observed that climate variables significantly improved (i.e. deltaAIC << 3) model performance beyond both null models for both drought and insects and that the range of P50 significantly improved drought mortality models, but not insect models.

We assessed model performance with cross-validation and used two primary metrics that reflect the performance of different parts of the hurdle model. We first tested for spatial autocorrelation using Moran’s I for each forest type and each of the drought and insect mortality models. For forest types and mortality models where significant autocorrelation was detected, we used a comparison of Moran’s I by distance bin, using the ‘correlog’ function in the pgirmess package in R (68), to determine the autocorrelation length. We set the spatial autocorrelation length as the midpoint between the last significant bin and the first non-significant bin. We then conducted spatial hold-out cross-validation (69) for each forest type and mortality model, whereby one grid cell was held out from model training and a spatial buffer around that grid cell equal to the autocorrelation length was also removed from model training. The model was then fit on the remaining data and used to predict the hold-out grid cell, and this was repeated 1000-fold for each forest type and mortality model.

Similar to the fire model, we examined model performance using cross-validated area under the receiver operating curve (AUC) for the binary component of the hurdle model and the non-zero-value *R^2^* for the beta-regression part of the hurdle model (Figure S5). We also compared the spatial correlations of observed and modeled mortality by aggregating across all modeled forest types via the weighted mean of the live basal area at t0 in each 0.25 degree grid cell. To calculate the cross-validated spatial correlation across the entire US, we randomly sampled the cross-validated predictions for each forest type with replacement, repeated this aggregation across forest types, and then repeated that process 1000 times (Figure S5).

Finally, we imposed one further set of criteria on drought and insect models to incorporate climate-dependence only where justified based on model performance. For all final model-based analyses (i.e. Figure 1D, 1F, Figures 2–4), we identified forest types where cross-validated AUC was greater than 0.6 and the forest type had >20 grid cells with mortality observed in the historical record, based on a recent analysis of stability and information criteria in regression models (70). This led to risks being modeled with climate variables and projected for 30 forest types in the insect models and 61 forest types in the drought models out of 112 possible forest types. For all forest types that did not meet these criteria, we projected mortality simply as the mean of historical mortality for that forest type, and thus set all future mortality to that value. We note that this is a very conservative decision and is very likely to underestimate future risks.

We performed a sensitivity analysis to ensure that our mortality models were robust to consideration of design weights that are assessed for FIA plots. FIA plots are often assigned weights based on strata and estimation units for given inventories. Following previous studies (61), we used the rFIA package (31) to extract weights for all our conditions and constructed both drought and insect mortality models with and without weighting. We found very strong agreement between the two approaches in both modeled mortality patterns (*R^2^*=0.95, *R^2^*=0.98, respectively; p<<0.00001 for both) and model coefficients (*R^2^*=0.88, *R^2^*=0.90, respectively; p<<0.00001 for both), indicating that our approach was robust to weighting decisions (Figure S6). We further checked to ensure that mortality driven by invasive species was not driving our mortality model patterns by verifying that our dominant and high-sensitivity forest types were not those known to be widely impacted by invasive species.

We further performed two evaluations of our mortality models against independent metrics or datasets. For the drought mortality models, we compared our observed mortality rates by species against the recent “Forest Stability Index” for eight major western US forest species (61) and observed a strong relationship (*R^2^*=0.72; Figure S2). For the insect mortality models, we compared our observed and modeled spatial patterns to an independent dataset of Aerial Detection Surveys done by the U.S. Forest Service to map bark beetle-driven mortality across the US (71). Despite large differences in types of dataset (e.g. aerial versus plot; “bark beetle-driven” mortality versus “insect-driven” plot agent codes) and spatial scales, we found strong agreement between our models and that independent dataset (AUC=0.79 and *R^2^*=0.29 comparing our modeled mortality and ADS observed mortality; Figure S3).

For future CMIP6 projections, we used the same climate variables as chosen by the final model selection analysis and projected drought and insect risk (i.e. % basal area killed per year) over each decade from 1950-2100 in different climate models and scenarios. Future decadal climate variables were z-scored against three decadal baselines (1990-1999; 2000-2009; 2010-2019) and the ensemble mean was taken across these baselines for each climate model and decade. All modeled and projection maps (e.g. Figure 1D/F, Figure 3) were made on all conditions in FIA, regardless of number of censuses in the historical record, to cover all US forests, aggregated to 0.25 degrees by forest type, and then aggregated across forest types as described above. For future projections, we used a constant stand age and P50 functional trait based on the 2000-2018 historical values due to uncertainties about future forest dynamics and composition. This is an assumption and uncertainty, but a full exploration of stand age dynamics, species distribution and composition shifts, and demography is beyond the scope of this current analysis. We note that future projections of hydrological variables included an empirical correction factor to account for CO2-related water savings effects from declines in stomatal conductance and thus our results at least partially account for CO2 as a potential ameliorating factor in mortality rates (46, 72). We verified that the mortality models predict reasonable mortality patterns and magnitudes when driven by CMIP6 historical model output (Figure 2), which gives confidence that future risks are due to climate change and not differences between the training TerraClimate data and CMIP6 data.

First-order estimates of 100-year integrated “reversal” (i.e. disturbance event that likely leads to a loss of biomass in a forest) risk were computed over the 2000-2099 period. We calculated these for the fires model based on the direct burn area historical data (2000-2018 average) and for SSP 5-8.5 based on the fractional change in each year compared to the 2000-2018 average. For insects, we calculated the historical reversal risk by tabulating the number of FIA conditions that lost >10% of biomass between the second and first measurement and had >0.5 fraction of mortality due to insect agent codes. For drought, we took a strongly conservative approach and tabulated conditions that lost >10% biomass after screening out conditions that had any documented agent codes of insects, disease, fire or vegetation or any disturbance codes of insect, human, weather, cutting, or any other human treatments. Furthermore, all drought reversal conditions had to have experienced PDSI minimum values of < −2 during the census interval. For all three disturbances, we used the multi-model mean to estimate a % change in historical risk in SSP 585 and simulated 1000 iterations (i.e. draws from a binomial distribution) of 100-year integrated risks with the given annual (fire) or (drought and insect) decadal risk probabilities.

### Open source software and data

We performed all analysis using open source software and publicly available data. Our team used the Pangeo cloud environment (73) to deploy a wide range of open source software packages in Python and R. Packages included: Pandas (74), Xarray (75); Matplotlib (76); NumPy (77); Seaborn (78); Jupyter (79); scikit-learn (55); Dask (80); GeoPandas (81); Zarr (82); RasterIO (88), MASS (73), betareg (62), pgirmess (68), raster (85), and ape (84). The source code to reproduce our analysis is available in https://doi.org/10.5281/zenodo.4741329. Archival versions of this project’s data products are available in https://doi.org/10.5281/zenodo.4741333.

## Acknowledgements and Funding

We thank H Stanke and J Shaw for insights and assistance with FIA data analysis and processing. Any errors are the authors’ sole responsibility. CarbonPlan acknowledges support from Microsoft AI for Earth. WRLA acknowledges support from the David and Lucille Packard Foundation, US National Science Foundation grants 1714972, 1802880 and 2003017, and USDA National Institute of Food and Agriculture, Agricultural and Food Research Initiative Competitive Programme, Ecosystem Services and Agro-Ecosystem Management, grant no. 2018-67019-27850. ATT acknowledges funding from the NSF Grant 2003205, the USDA National Institute of Food and Agriculture, Agricultural and Food Research Initiative Competitive Programme Grant No. 2018-67012-31496 and the University of California Laboratory Fees Research Program Award No. LFR-20-652467.

## Supplementary Information

### Here we provide some additional information around our fire analysis, as well as eight additional figures related to all of our analyses

The fitted model predicts patterns of historical fire with a cross-validated AUC of 0.89 and *R^2^* of 0.067. Comparing spatial patterns of observed burn area and model predictions for the historical period 1984 through 2018 reveals moderate agreement (Figure 1a) with a cross-validated spatial *R^2^* of 0.057. The model notably underpredicts burn area in coastal Northern California and Southern Oregon, as well the Tetons in northwestern Wyoming. Another key difference is the model’s weak and spatially diffuse level of elevated risk across a large swath of the Western US, for example in the Northern Rocky Mountains and central Oklahoma. This behavior is generally expected as the model, by design, includes no stochastic element of triggers like lightning or human activity and is aiming to capture long-term fire risk, rather than individual fire events. As a result, the model predicts broad conditions over large spatial areas that could lead to fires, as opposed to exactly predicting triggered fire events.

The model robustly captures key temporal trends in the MTBS fire record (seasonal cross-validated *R^2^*: 0.92; inter-annual cross-validated *R^2^*: 0.62). Comparing observed interannual variability (Figure S8), the model captures periods of high fire (e.g. 2005-2006, 2011-2012) and receding in the troughs (e.g. 2004, 2009-2010, 2014). The model also captures the overall increase in fire risk over the 35 year historical record. The model reproduces the average monthly seasonal cycle (middle panel) predicting higher fire risk in the summer, followed by lulls in the winter. The full monthly time series is detailed in the bottom panel. Notably, the model underpredicts the highest, most extreme summertime spikes in fire risk (e.g. 2007). As is common in time series modeling, the model effectively reproduces the mean behavior and thus tends to underestimate peaks with an elevated baseline.

The model also captures the spatial distribution of fire as it evolves throughout the fire season, as seen in monthly maps of burn area predictions (Figure S7). The model predicts the springtime fire season in the Southwest and Southeast, followed by the comparatively stronger summer and fall fire risk in the Western US. As in Figure 1, the model predicts a larger, more diffuse spatial extent with lower peaks compared to the observed record. This aligns with the design of the statistical model. As an example, peaks of high burn area in the Northern Rockies of Idaho and Montana in the observational record in July and August are modeled with a smoother extent of elevated risk across the entire region.

**Figure S1.**
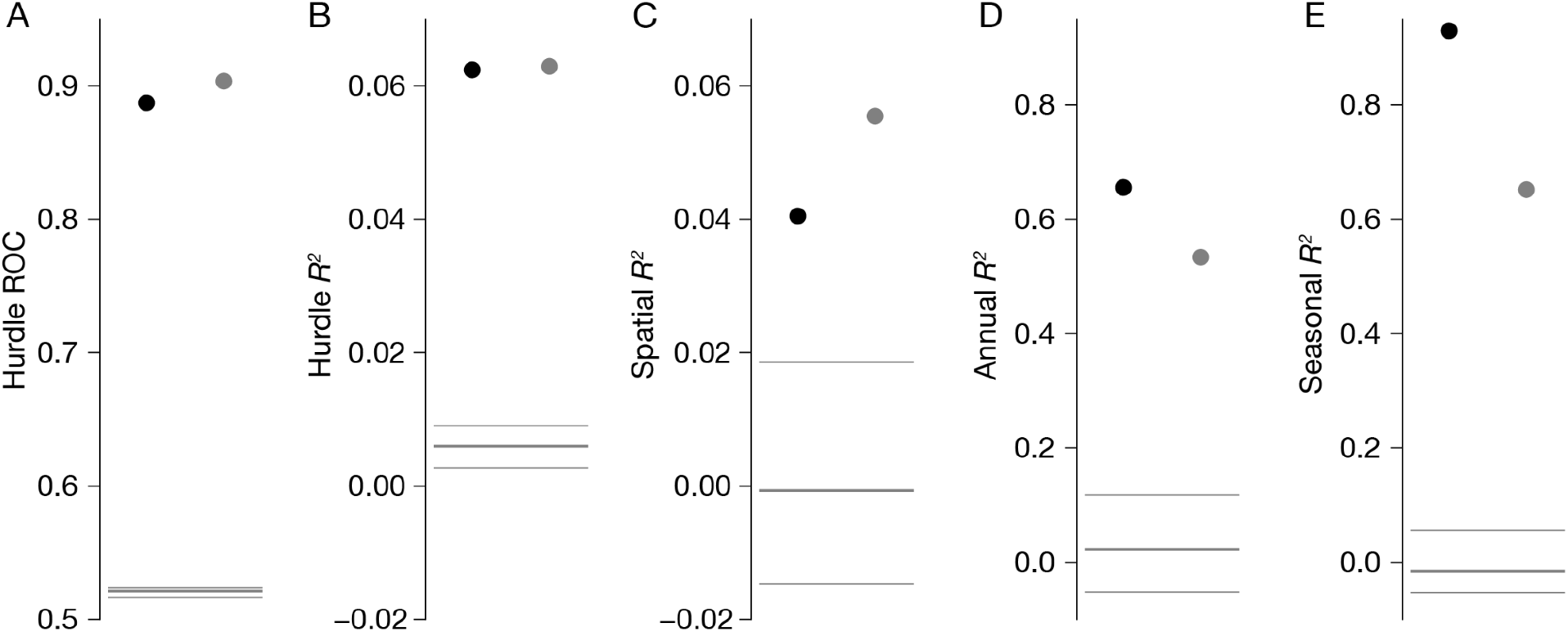
Cross-validation performance and shuffle statistics for the fire model. Each panel shows a single performance metric with two two different forms of cross-validation (black dot, split-halves; gray dot, extrapolation) compared to a null distribution generated by shuffling the relationship between training and testing (thick gray line, median; thin gray lines, minimum and maximum). The shuffle distribution provides an indication of the performance that might be expected due to fitting noise. For spatial, annual, and seasonal *R^2^* we allow for an arbitrary offset, so we separately computed the bias when cross-validating these predictions (median 2.4% bias for split-halves and −52% for extrapolation). The substantial negative bias for extrapolation suggests that our projected estimates of fire risk are, if anything, conservative.

**Figure S2.**
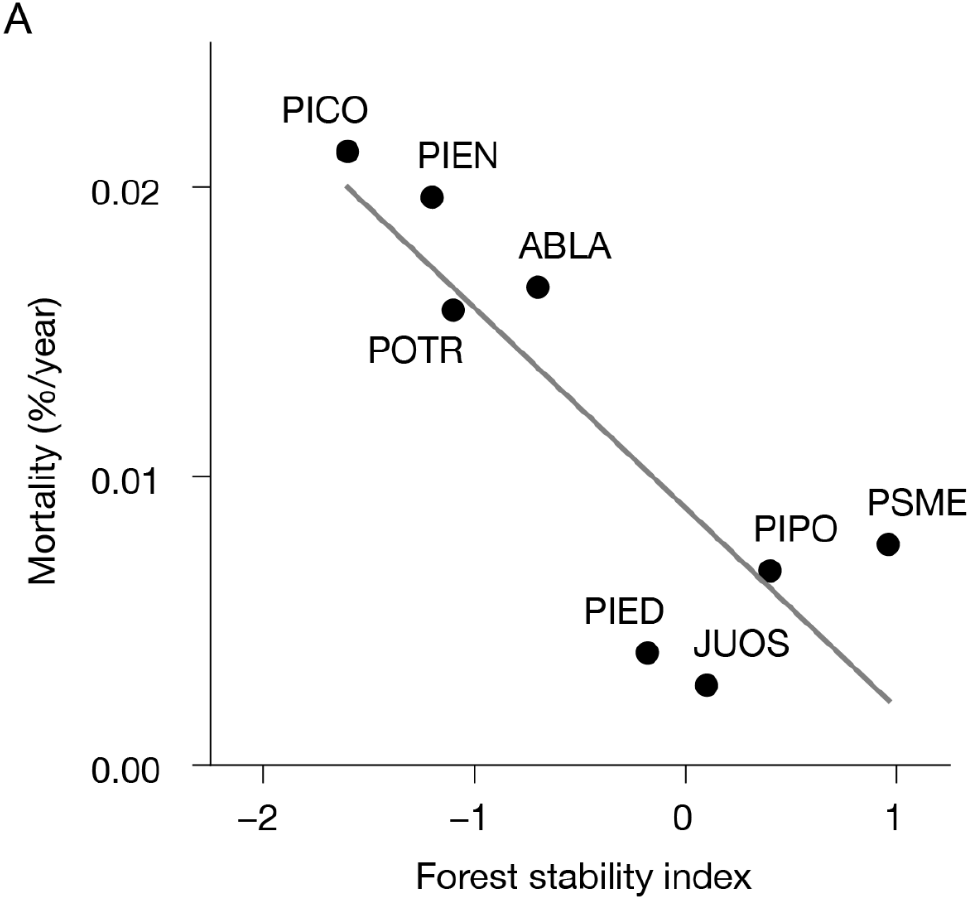
Comparison of forest type-level average mortality rates (% basal area killed / year) for major western US forest types from our data to a recently published “forest stability index” (FSI) (86) at a species level. Tree species are Pinus contorta (PICO; lodgepole pine forest type in this analysis), Picea engelmanii (PIEN; Engelmann spruce forest type), Populus tremuloides (POTR; aspen forest type), Abies lasiocarpa (ABLA; subalpine fir forest type), Pinus edulis (PIED; pinon-juniper woodland forest type), Juniperus osteosperma (JUOS; juniper woodland forest type), Pinus ponderosa (PIPO; ponderosa pine forest type), and Pseudotsuga menziesii (PSME; douglas fir forest type). Gray line shows a linear regression.

**Figure S3.**
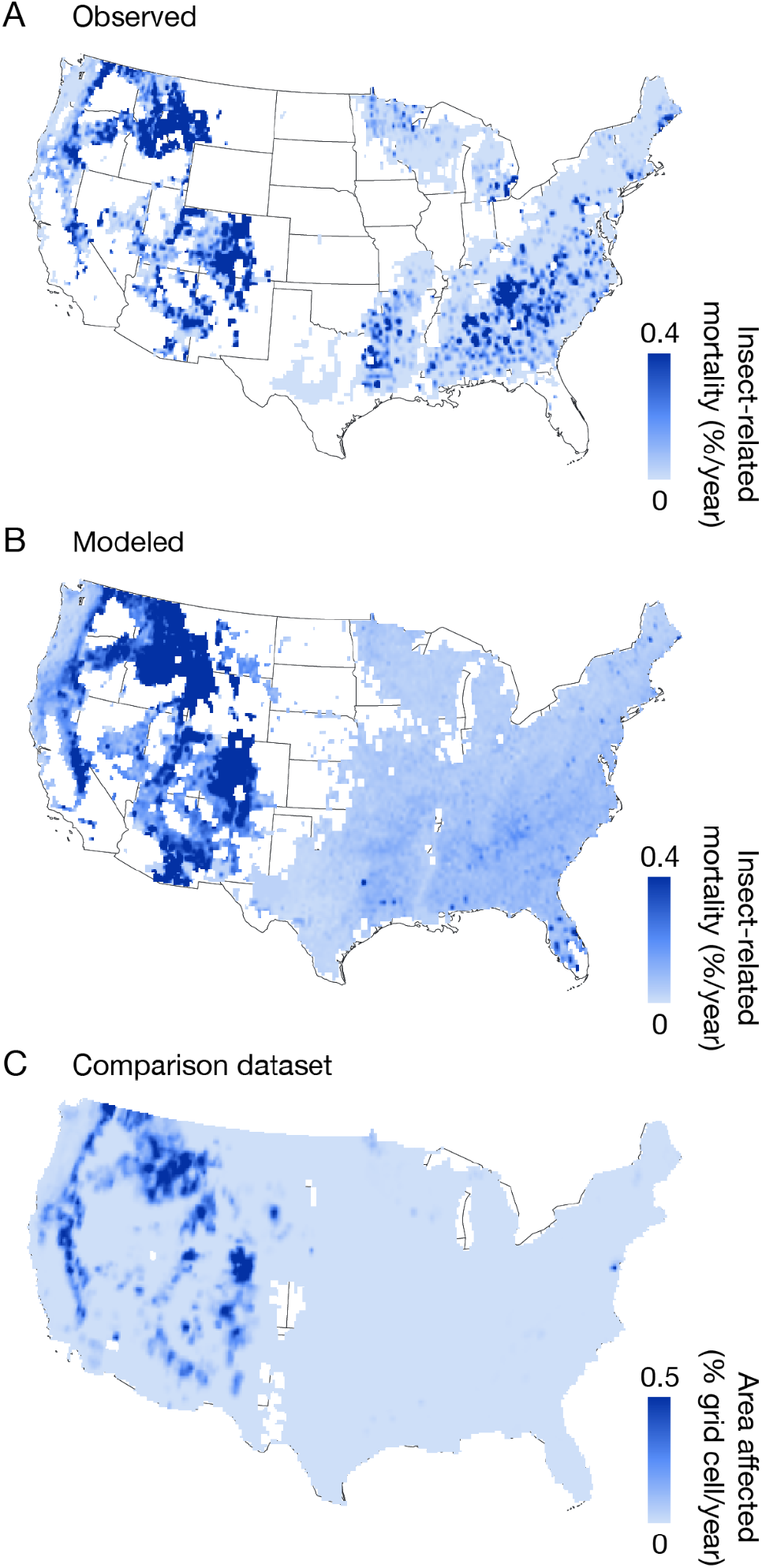
Comparison of FIA-derived observed (A) and modeled (B) insect-driven tree mortality (% basal area killed / year) to an independent bark-beetle tree mortality dataset (C) from aerial detection surveys (% of grid cell area affected / year) (71).

**Figure S4.**
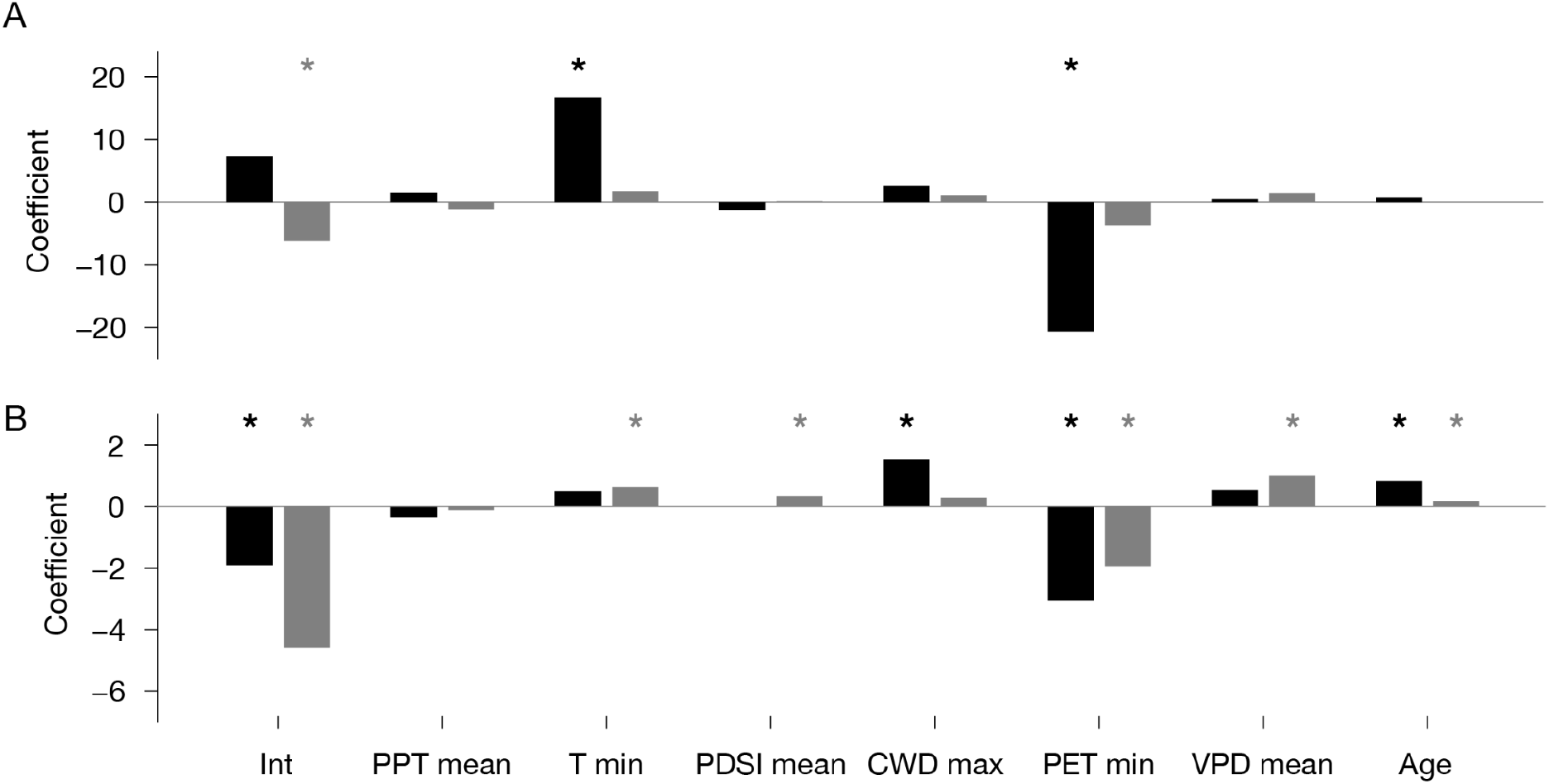
Insect-driven tree mortality climate sensitivities are consistent with previous studies finding connections between temperature and drought stress and insect-driven tree mortality *(36, 87)*. Model coefficients for the binomial (black) and beta (gray) portions of the hurdle model are shown for two example forest types, limber pine (A) and lodgepole pine (B).

**Figure S5.**
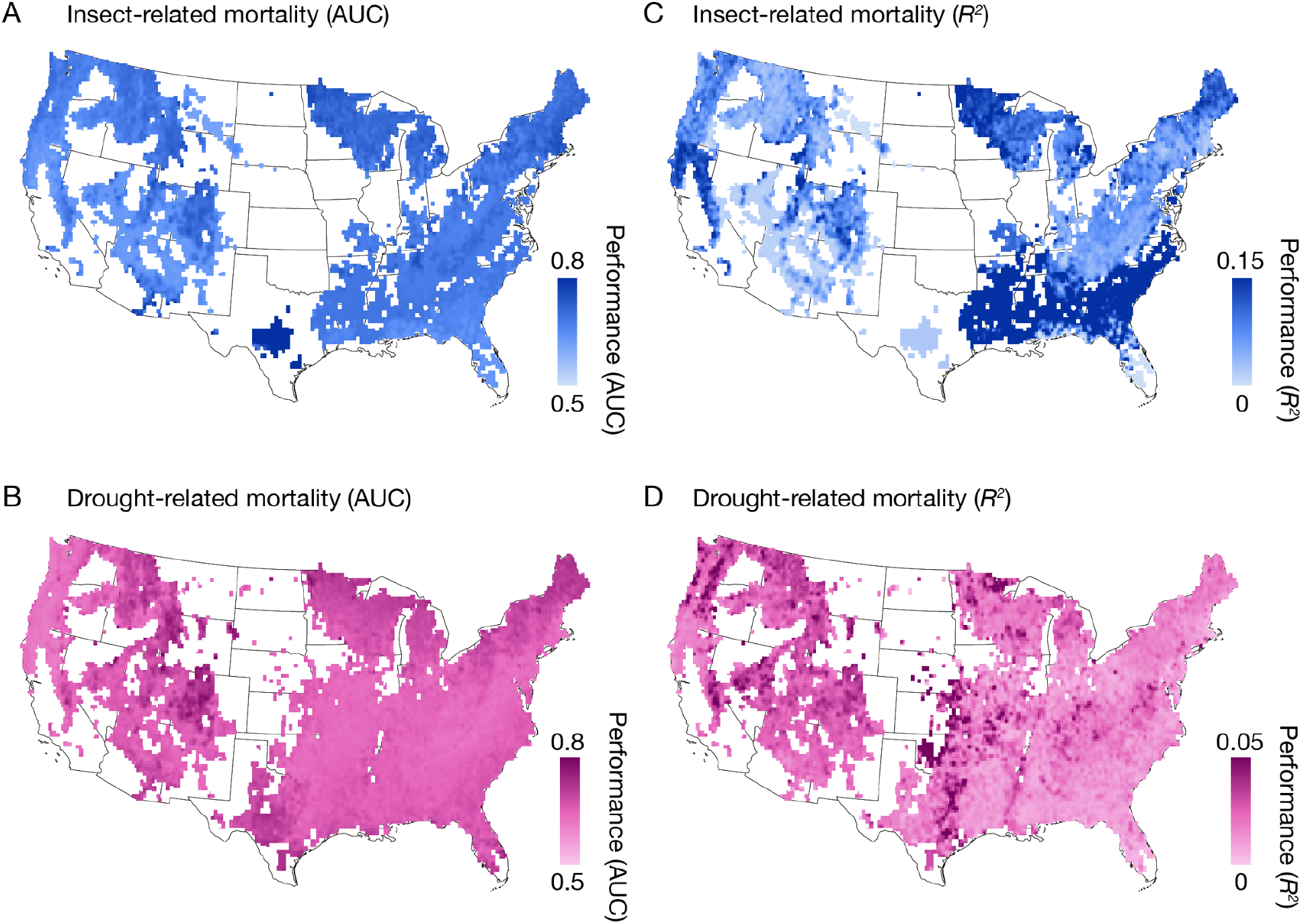
Drought and insect model performance. Cross-validated area under receiver operator curve (AUC) for insect (A) and drought (B) models. Cross-validated *R^2^* values for grid cells with non-zero mortality for insect (C) and drought (D) models. We separately performed a spatial cross-validation yield *R^2^* values of 0.18 (90% CI: 0.16 to 0.20) for drought models and 0.31 (90% CI: 0.23 to 0.34) for insect models.

**Figure S6.**
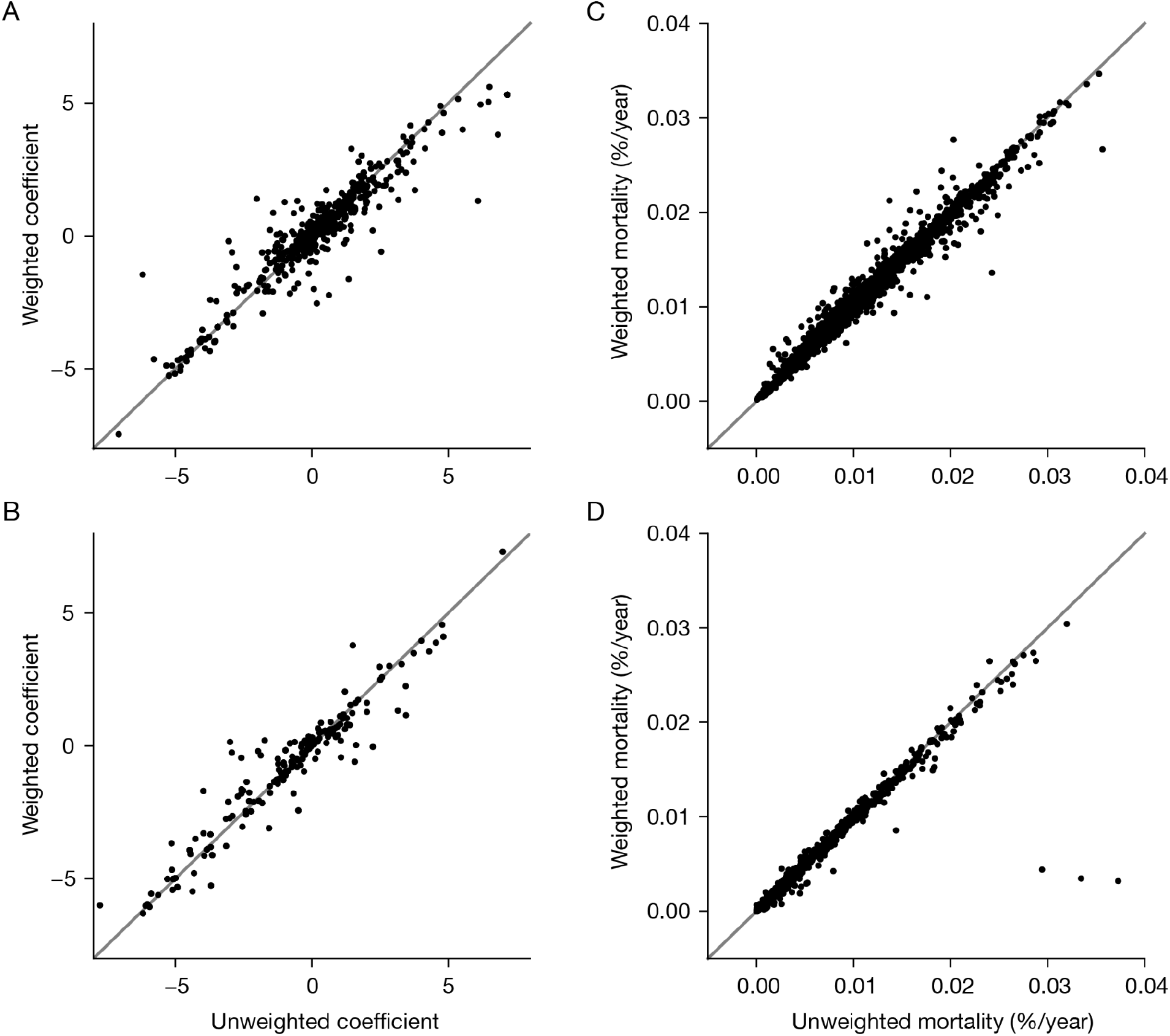
Mortality models are highly consistent across Forest Inventory and Analysis (FIA) stratum weighting methods. (A, B) Continental drought and insect coefficients from regression models with or without including FIA weights (*R^2^*=0.90, *R^2^*=0.88, p<<0.00001 for both). (C, D) Continental drought and insect mortality predictions from regression models with or without including FIA weights (*R^2^*=0.98, *R^2^*=0.95, p<<0.00001 for both).

**Figure S7.**
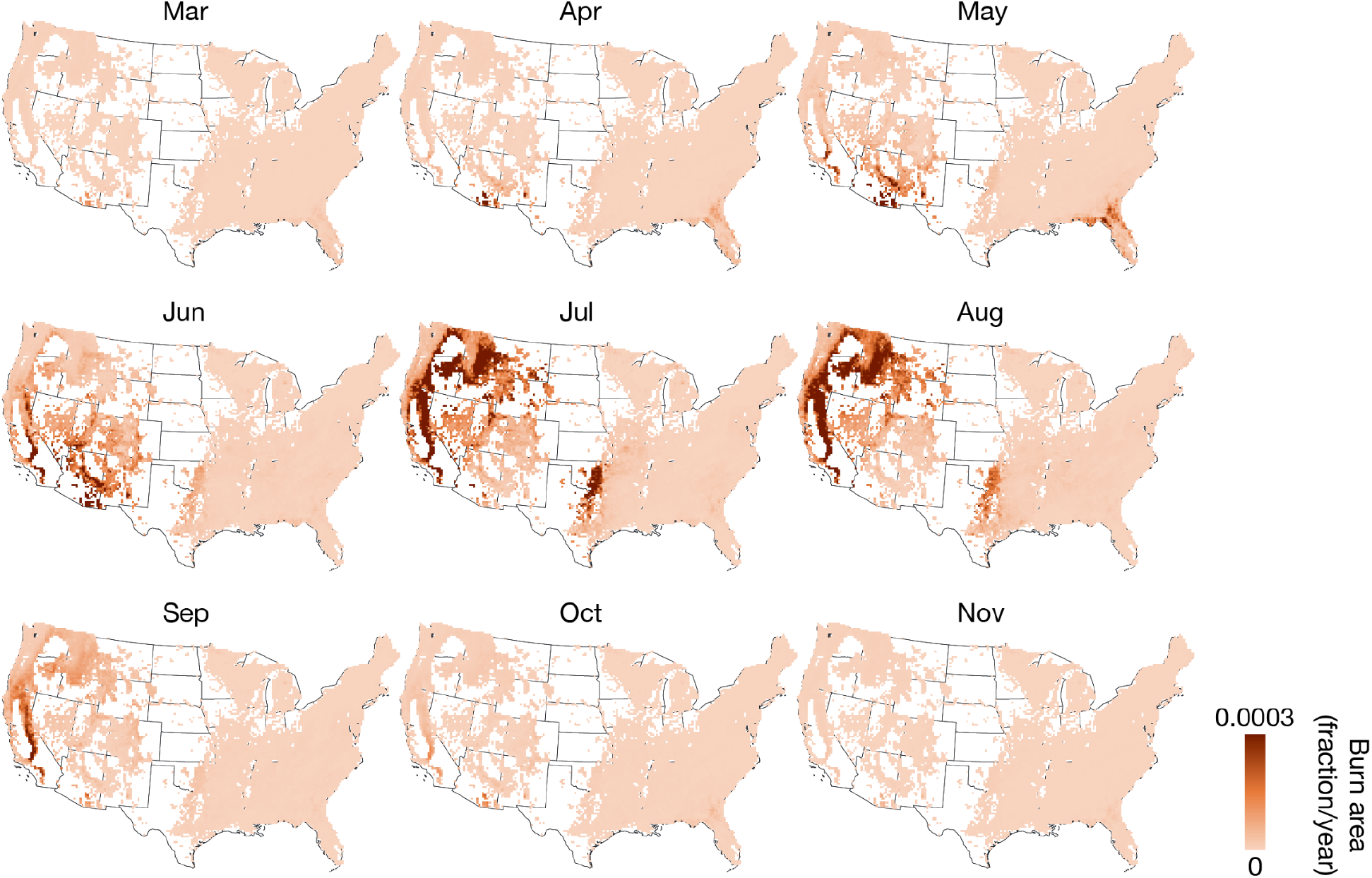
Seasonal patterns of burn area from model predictions, averaged over the 1984-2018 historical period. The model captures the observed evolution of spatial burn area patterns throughout the year. For example, modeled burn area in the Southeastern US peaks in the Spring, followed by increases in the intermountain west in the summer. Modeled burn area in Autumn is predominantly within California.

**Figure S8.**
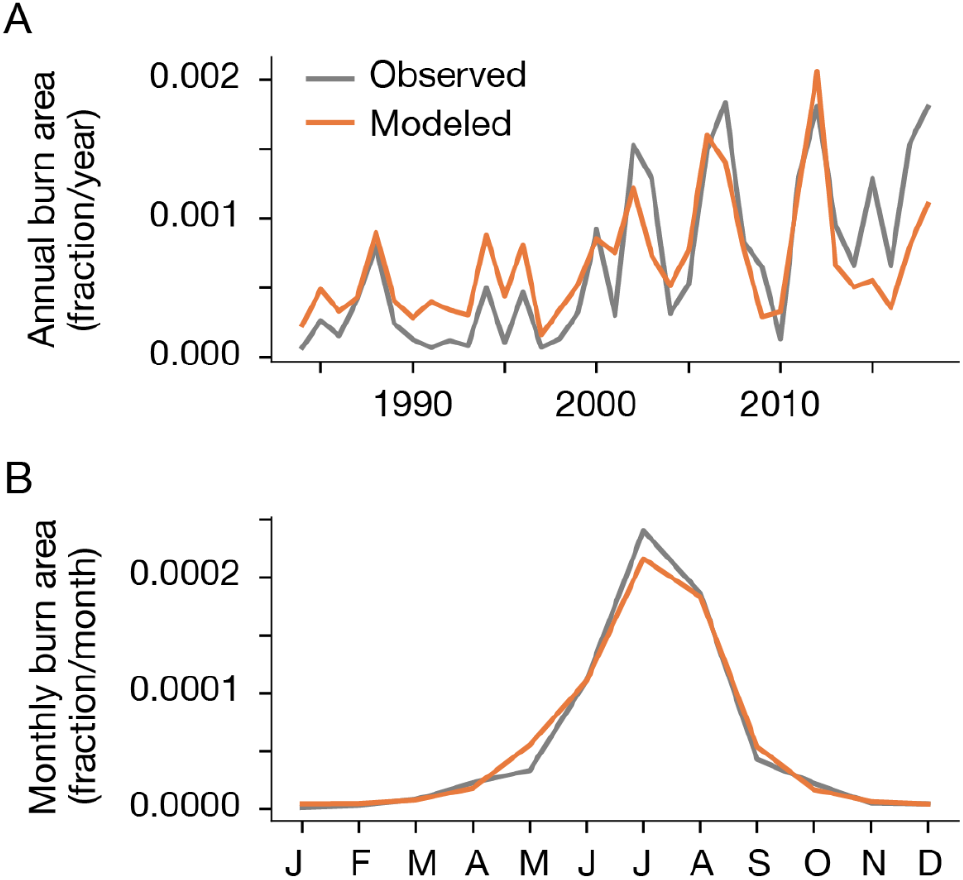
Temporal trends in observed and modeled burn area averaged over the continental United States for the historical period 1984-2018. As seen in the annual average (A) the model captures interannual variability as well as increases over the later part of the record. The model also captures the seasonal pattern (B) peaking in summer.

## Notes

https://doi.org/10.5281/zenodo.4741333

https://doi.org/10.5281/zenodo.4741329

### Competing Interest Statement

The authors have declared no competing interest.

## References

1. G. B. Bonan, Forests and climate change: Forcings, feedbacks, and the climate benefits of forests. Science. 320, 1444–1449 (2008).

2. Y. Pan, R. A. Birdsey, J. Fang, R. Houghton, P. E. Kauppi, W. A. Kurz, O. L. Phillips, A. Shvidenko, S. L. Lewis, J. G. Canadell, P. Ciais, R. B. Jackson, S. W. Pacala, A. D. McGuire, S. Piao, A. Rautiainen, S. Sitch, D. Hayes, A Large and Persistent Carbon Sink in the World’s Forests. Science. 333, 988–993 (2011).

3. B. W. Griscom, J. Adams, P. W. Ellis, R. A. Houghton, G. Lomax, D. A. Miteva, W. H. Schlesinger, D. Shoch, J. V. Siikamäki, P. Smith, Natural climate solutions. Proc. Natl.Acad. Sci. 114, 11645–11650 (2017).

4. J. E. Fargione, S. Bassett, T. Boucher, S. D. Bridgham, R. T. Conant, S. C. Cook-Patton, P.W. Ellis, A. Falcucci, J. W. Fourqurean, T. Gopalakrishna, Natural climate solutions for the United States. Sci. Adv. 4, eaat1869 (2018).

5. S. Roe, C. Streck, M. Obersteiner, S. Frank, B. Griscom, L. Drouet, O. Fricko, M. Gusti, N. Harris, T. Hasegawa, Contribution of the land sector to a 1.5° C world. Nat. Clim. Change, 1–12 (2019).

6. S. C. Cook-Patton, S. M. Leavitt, D. Gibbs, N. L. Harris, K. Lister, K. J. Anderson-Teixeira, R. D. Briggs, R. L. Chazdon, T. W. Crowther, P. W. Ellis, Mapping carbon accumulation potential from global natural forest regrowth. Nature. 585, 545–550 (2020).

7. M. D. Hurteau, B. A. Hungate, G. W. Koch, Accounting for risk in valuing forest carbon offsets. Carbon Balance Manag. 4, 1 (2009).

8. T. Ruseva, E. Marland, C. Szymanski, J. Hoyle, G. Marland, T. Kowalczyk, Additionality and permanence standards in California’s Forest Offset Protocol: A review of project and program level implications. J. Environ. Manage. 198, 277–288 (2017).

9. D. Archer, M. Eby, V. Brovkin, A. Ridgwell, L. Cao, U. Mikolajewicz, K. Caldeira, K. Matsumoto, G. Munhoven, A. Montenegro, Atmospheric lifetime of fossil fuel carbon dioxide. Annu. Rev. Earth Planet. Sci. 37(2009).

10. P. Friedlingstein, M. Meinshausen, V. K. Arora, C. D. Jones, A. Anav, S. K. Liddicoat, R. Knutti, Uncertainties in CMIP5 Climate Projections due to Carbon Cycle Feedbacks. J.Clim. 27(2014).

11. T. J. Brodribb, J. Powers, H. Cochard, B. Choat, Hanging by a thread? Forests and drought. Science. 368, 261–266 (2020).

12. R. Seidl, D. Thom, M. Kautz, D. Martin-Benito, M. Peltoniemi, G. Vacchiano, J. Wild, D. Ascoli, M. Petr, J. Honkaniemi, Forest disturbances under climate change. Nat. Clim.Change. 7, 395 (2017).

13. D. Baldocchi, J. Penuelas, The physics and ecology of mining carbon dioxide from the atmosphere by ecosystems. Glob. Change Biol. 25, 1191–1197 (2019).

14. T. G. Holland, W. Stewart, M. D. Potts, Source or Sink? A comparison of Landfire-and FIA-based estimates of change in aboveground live tree carbon in California’s forests. Environ. Res. Lett. 14, 074008 (2019).

15. W. A. Kurz, C. C. Dymond, G. Stinson, G. J. Rampley, E. T. Neilson, A. L. Carroll, T. Ebata, L. Safranyik, Mountain pine beetle and forest carbon feedback to climate change. Nature. 452, 987–990 (2008).

16. W. A. Kurz, G. Stinson, G. J. Rampley, C. C. Dymond, E. T. Neilson, Risk of natural disturbances makes future contribution of Canada’s forests to the global carbon cycle highly uncertain. Proc. Natl. Acad. Sci. 105, 1551–1555 (2008).

17. B. J. Bentz, J. Régnière, C. J. Fettig, E. M. Hansen, J. L. Hayes, J. A. Hicke, R. G. Kelsey, J. F. Negrón, S. J. Seybold, Climate Change and Bark Beetles of the Western United States and Canada: Direct and Indirect Effects. BioScience. 60, 602–613 (2010).

18. A. P. Williams, C. D. Allen, A. K. Macalady, D. Griffin, C. A. Woodhouse, D. M. Meko, T. W. Swetnam, S. A. Rauscher, R. Seager, H. D. Grissino-Mayer, J. S. Dean, E. R. Cook, C. Gangodagamage, M. Cai, N. G. McDowell, Temperature as a potent driver of regional forest drought stress and tree mortality. Nat. Clim Change. 3, 292–297 (2013).

19. J. A. Hicke, C. D. Allen, A. R. Desai, M. C. Dietze, R. J. Hall, D. M. Kashian, D. Moore, K.F. Raffa, R. N. Sturrock, J. Vogelmann, Effects of biotic disturbances on forest carbon cycling in the United States and Canada. Glob. Change Biol. 18, 7–34 (2012).

20. R. Barbero, J. T. Abatzoglou, N. K. Larkin, C. A. Kolden, B. Stocks, Climate change presents increased potential for very large fires in the contiguous United States. Int. J.Wildland Fire. 24, 892–899 (2015).

21. P. C. Buotte, S. Levis, B. E. Law, T. W. Hudiburg, D. E. Rupp, J. J. Kent, Near-future forest vulnerability to drought and fire varies across the western United States. Glob. Change Biol. 25, 290–303 (2019).

22. J. A. Wang, A. Baccini, M. Farina, J. T. Randerson, M. A. Friedl, Disturbance suppresses the aboveground carbon sink in North American boreal forests. Nat. Clim. Change (2021),doi:10.1038/s41558-021-01027-4.

23. X. J. Walker, J. L. Baltzer, S. G. Cumming, N. J. Day, C. Ebert, S. Goetz, J. F. Johnstone, S. Potter, B. M. Rogers, E. A. G. Schuur, M. R. Turetsky, M. C. Mack, Increasing wildfires threaten historic carbon sink of boreal forest soils. Nature. 572, 520–523 (2019).

24. M. Reichstein, M. Bahn, P. Ciais, D. Frank, M. D. Mahecha, S. I. Seneviratne, J. Zscheischler, C. Beer, N. Buchmann, D. C. Frank, others, Climate extremes and the carbon cycle. Nature. 500, 287–295 (2013).

25. A. P. Williams, J. T. Abatzoglou, Recent advances and remaining uncertainties in resolving past and future climate effects on global fire activity. Curr. Clim. Change Rep. 2, 1–14(2016).

26. J. S. Clark, L. Iverson, C. W. Woodall, C. D. Allen, D. M. Bell, D. C. Bragg, A. W. D’amato, F. W. Davis, M. H. Hersh, I. Ibanez, others, The impacts of increasing drought on forest dynamics, structure, and biodiversity in the United States. Glob. Change Biol. (2016)(available at http://onlinelibrary.wiley.com/doi/10.1111/gcb.13160/pdf).

27. W. R. Anderegg, A. T. Trugman, G. Badgley, C. M. Anderson, A. Bartuska, P. Ciais, D. Cullenward, C. B. Field, J. Freeman, S. J. Goetz, Climate-driven risks to the climate mitigation potential of forests. Science. 368(2020).

28. J. Lecina-Diaz, J. Martínez-Vilalta, A. Alvarez, M. Banqué, J. Birkmann, D. Feldmeyer, J. Vayreda, J. Retana, Characterizing forest vulnerability and risk to climate-change hazards. Front. Ecol. Environ. 19, 126–133 (2021).

29. M. D. Hurteau, B. A. Hungate, G. W. Koch, M. P. North, G. R. Smith, Aligning ecology and markets in the forest carbon cycle. Front. Ecol. Environ. 11, 37–42 (2013).

30. R. Barbero, J. T. Abatzoglou, E. A. Steel, N. K Larkin, Modeling very large-fire occurrences over the continental United States from weather and climate forcing. Environ. Res. Lett. 9, 124009 (2014).

31. H. Stanke, A. O. Finley, A. S. Weed, B. F. Walters, G. M. Domke, rFIA: An R package for estimation of forest attributes with the US Forest Inventory and Analysis database. Environ. Model. Softw. 127, 104664 (2020).

32. A. T. Trugman, L. D. Anderegg, J. D. Shaw, W. R. Anderegg, Trait velocities reveal that mortality has driven widespread coordinated shifts in forest hydraulic trait composition. Proc. Natl. Acad. Sci. 117, 8532–8538 (2020).

33. W. R. Anderegg, T. Klein, M. Bartlett, L. Sack, A. F. Pellegrini, B. Choat, S. Jansen, Meta-analysis reveals that hydraulic traits explain cross-species patterns of drought-induced tree mortality across the globe. Proc. Natl. Acad. Sci., 201525678 (2016).

34. A. J. Meddens, J. A. Hicke, A. K. Macalady, P. C. Buotte, T. R. Cowles, C. D. Allen, Patterns and causes of observed piñon pine mortality in the southwestern United States. New Phytol. 206, 91–97 (2015).

35. E. P. Creeden, J. A. Hicke, P. C. Buotte, Climate, weather, and recent mountain pine beetle outbreaks in the western United States. For. Ecol. Manag. 312, 239–251 (2014).

36. B. J. Bentz, J. Régnière, C. J. Fettig, E. M. Hansen, J. L. Hayes, J. A. Hicke, R. G. Kelsey, J. F. Negrón, S. J. Seybold, Climate Change and Bark Beetles of the Western United States and Canada: Direct and Indirect Effects. BioScience. 60, 602–613 (2010).

37. B. E. McNellis, A. M. Smith, A. T. Hudak, E. K. Strand, Tree mortality in western US forests forecasted using forest inventory and Random Forest classification. Ecosphere. 12, e03419 (2021).

38. A. S. Weed, M. P. Ayres, J. Hicke, Consequences of climate change for biotic disturbances in North American forests. Ecol. Monogr. (2013).

39. A. M. Smith, C. A. Kolden, T. B. Paveglio, M. A. Cochrane, D. M. Bowman, M. A. Moritz, A. D. Kliskey, L. Alessa, A. T. Hudak, C. M. Hoffman, The science of firescapes: achieving fire-resilient communities. Bioscience. 66, 130–146 (2016).

40. R. A. Fisher, C. D. Koven, W. R. Anderegg, B. O. Christoffersen, M. C. Dietze, C. E. Farrior, J. A. Holm, G. C. Hurtt, R. G. Knox, P. J. Lawrence, Vegetation demographics in Earth System Models: A review of progress and priorities. Glob. Change Biol. 24, 35–54(2018).

41. H. Bugmann, R. Seidl, F. Hartig, F. Bohn, J. Br\uuna, M. Cailleret, L. François, J. Heinke, A.-J. Henrot, T. Hickler, Tree mortality submodels drive simulated long-term forest dynamics: assessing 15 models from the stand to global scale. Ecosphere. 10, e02616(2019).

42. T. A. Pugh, A. Arneth, M. Kautz, B. Poulter, B. Smith, Important role of forest disturbances in the global biomass turnover and carbon sinks. Nat. Geosci. 12, 730–735 (2019).

43. J. Eidenshink, B. Schwind, K. Brewer, Z.-L. Zhu, B. Quayle, S. Howard, A Project for Monitoring Trends in Burn Severity. Fire Ecol. 3, 3–21 (2007).

44. B. Ruefenacht, M. V. Finco, M. D. Nelson, R. Czaplewski, E. H. Helmer, J. A. Blackard, G.R. Holden, A. J. Lister, D. Salajanu, D. Weyermann, K. Winterberger, Conterminous U.S.*and Alaska Forest Type Mapping Using Forest Inventory and Analysis Data*. Photogramm.Eng. Remote Sens. 74, 1379–1388 (2008).

45. J. Dewitz, National Land Cover Dataset (NLCD) 2016 Products (2019), doi:10.5066/P96HHBIE.

46. J. T. Abatzoglou, S. Z. Dobrowski, S. A. Parks, K. C. Hegewisch, TerraClimate, a high-resolution global dataset of monthly climate and climatic water balance from 1958-2015. Sci. Data. 5, 170191 (2018).

47. V. Eyring, S. Bony, G. A. Meehl, C. A. Senior, B. Stevens, R. J. Stouffer, K. E. Taylor, Overview of the Coupled Model Intercomparison Project Phase 6 (CMIP6) experimental design and organization. Geosci. Model Dev. 9, 1937–1958 (2016).

48. S. R. Sobie, T. Q. Murdock, High-Resolution Statistical Downscaling in Southwestern British Columbia. J. Appl. Meteorol. Climatol. 56, 1625–1641 (2017).

49. H. Li, J. Sheffield, E. F. Wood, Bias correction of monthly precipitation and temperature fields from Intergovernmental Panel on Climate Change AR4 models using equidistant quantile matching. J. Geophys. Res. 115, D10101 (2010).

50. A. W. Wood, L. R. Leung, V. Sridhar, D. P. Lettenmaier, Hydrologic Implications of Dynamical and Statistical Approaches to Downscaling Climate Model Outputs. Clim.Change. 62, 189–216 (2004).

51. D. Maraun, Bias Correction, Quantile Mapping, and Downscaling: Revisiting the Inflation Issue. J. Clim. 26, 2137–2143 (2013).

52. A. M. K. Stoner, K. Hayhoe, X. Yang, D. J. Wuebbles, An asynchronous regional regression model for statistical downscaling of daily climate variables: ASYNCHRONOUS REGRESSION MODEL FOR STATISTICAL CLIMATE DOWNSCALING. Int. J. Climatol. 33, 2473–2494 (2013).

53. Adams, James, climate_indices, an open source Python library providing reference implementations of commonly used climate indices (2017;https://github.com/monocongo/climate_indices).

54. J. G. Cragg, Some Statistical Models for Limited Dependent Variables with Application to the Demand for Durable Goods. Econometrica. 39, 829 (1971).

55. F. Pedregosa, G. Varoquaux, A. Gramfort, V. Michel, B. Thirion, O. Grisel, M. Blondel, P. Prettenhofer, R. Weiss, V. Dubourg, J. Vanderplas, A. Passos, D. Cournapeau, M. Brucher, M. Perrot, E. Duchesnay, G. Louppe, Scikit-learn: Machine Learning in Python. J. Mach.Learn. Res. 12(2012).

56. D. L. Swain, A. Shorter, Sharper Rainy Season Amplifies California Wildfire Risk. Geophys.Res. Lett. 48(2021), doi:10.1029/2021GL092843.

57. W. R. Anderegg, J. A. Hicke, R. A. Fisher, C. D. Allen, J. Aukema, B. Bentz, S. Hood, J. W. Lichstein, A. K. Macalady, N. McDowell, others, Tree mortality from drought, insects, and their interactions in a changing climate. New Phytol. 208, 674–683 (2015).

58. J. D. Shaw, B. E. Steed, L. T. DeBlander, Forest Inventory and Analysis (FIA) Annual Inventory Answers the Question: What Is Happening to Pinyon-Juniper Woodlands? J. For. 103, 280–285 (2005).

59. R. A. Hember, W. A. Kurz, N. C. Coops, Relationships between individual-tree mortality and water-balance variables indicate positive trends in water stress-induced tree mortality across North America. Glob. Change Biol. 23, 1691–1710 (2017).

60. W. R. Anderegg, A. T. Trugman, G. Badgley, A. G. Konings, J. Shaw, Divergent forest sensitivity to repeated extreme droughts. Nat. Clim. Change. 10, 1091–1095 (2020).

61. H. Stanke, A. O. Finley, G. M. Domke, A. S. Weed, D. W. MacFarlane, Over half of western United States’ most abundant tree species in decline. Nat. Commun. 12, 1–11 (2021).

62. A. Zeileis, F. Cribari-Neto, B. Grün, I. Kos-midis, Beta regression in R. J. Stat. Softw. 34, 1–24 (2010).

63. W. R. L. Anderegg, Alan Flint, Huang, Cho-ying, Flint, Lorraine, Berry, Joseph, Davis, Frank, Sperry, John, Field, Christopher, Tree mortality predicted from drought-induced vascular damage. Nat. Geosci. 8, 367–371 (2015).

64. C. Grossiord, T. N. Buckley, L. A. Cernusak, K. A. Novick, B. Poulter, R. T. Siegwolf, J. S. Sperry, N. G. McDowell, Plant responses to rising vapor pressure deficit. New Phytol. 226, 1550–1566 (2020).

65. A. Nardini, M. Battistuzzo, T. Savi, Shoot desiccation and hydraulic failure in temperate woody angiosperms during an extreme summer drought. New Phytol. 200, 322–329(2013).

66. M. Urli, A. J. Porté, H. Cochard, Y. Guengant, R. Burlett, S. Delzon, Xylem embolism threshold for catastrophic hydraulic failure in angiosperm trees. Tree Physiol. 33, 672–683(2013).

67. K. P. Burnham, D. R. Anderson, Multimodel inference understanding AIC and BIC in model selection. Sociol. Methods Res. 33, 261–304 (2004).

68. P. Giraudoux, M. P. Giraudoux, S. Mass, Package ‘pgirmess’ (2018).

69. P. Ploton, F. Mortier, M. Réjou-Méchain, N. Barbier, N. Picard, V. Rossi, C. Dormann, G. Cornu, G. Viennois, N. Bayol, A. Lyapustin, S. Gourlet-Fleury, R. Pélissier, Spatial validation reveals poor predictive performance of large-scale ecological mapping models. Nat. Commun. 11, 4540 (2020).

70. D. G. Jenkins, P. F. Quintana-Ascencio, A solution to minimum sample size for regressions. PloS One. 15, e0229345 (2020).

71. C. A. Williams, H. Gu, R. MacLean, J. G. Masek, G. J. Collatz, Disturbance and the carbon balance of US forests: A quantitative review of impacts from harvests, fires, insects, and droughts. Glob. Planet. Change. 143, 66–80 (2016).

72. J. S. Sperry, M. D. Venturas, H. N. Todd,A. T. Trugman, W. R. Anderegg, Y. Wang, X. Tai, The impact of rising CO2 and acclimation on the response of US forests to global warming. Proc. Natl. Acad. Sci. 116, 25734–25744 (2019).

73. N. H. Robinson, J. Hamman, R. Abernathey, Seven Principles for Effective Scientific Big-DataSystems. ArXiv190803356 Cs (2020) (available at http://arxiv.org/abs/1908.03356).

74. McKinney, Wes., in Proceedings of the 9th Python in Science Conference (SCIPY 2010)(2010), pp. 56–61.

75. S. Hoyer, J. J. Hamman, xarray: N-D labeled Arrays and Datasets in Python. J. Open Res.Softw. 5, 10 (2017).

76. J. D. Hunter, Matplotlib: A 2D Graphics Environment. Comput. Sci. Eng. 9, 90–95 (2007).

77. C. R. Harris, K. J. Millman, S. J. van der Walt, R. Gommers, P. Virtanen, D. Cournapeau, E. Wieser, J. Taylor, S. Berg, N. J. Smith, R. Kern, M. Picus, S. Hoyer, M. H. van Kerkwijk, M. Brett, A. Haldane, J. F. del Río, M. Wiebe, P. Peterson, P. Gérard-Marchant, K. Sheppard, T. Reddy, W. Weckesser, H. Abbasi, C. Gohlke, T. E. Oliphant, Array programming with NumPy. Nature. 585, 357–362 (2020).

78. M. Waskom, M. Gelbart, O. Botvinnik, J. Ostblom, P. Hobson, S. Lukauskas, D. C. Gemperline, T. Augspurger, Y. Halchenko, J. Warmenhoven, J. B. Cole, J. D. Ruiter, J. Vanderplas, S. Hoyer, C. Pye, A. Miles, Corban Swain, K. Meyer, M. Martin, P. Bachant, E. Quintero, G. Kunter, S. Villalba, Brian, C. Fitzgerald, C. Evans, M. L. Williams, D. O’Kane, T. Yarkoni, T. Brunner, mwaskom/seaborn: v0.11.1 (December 2020) (Zenodo, 2020;https://zenodo.org/record/4379347).

79. T. Kluyver, B. Ragan-Kelley, F. Pérez, B. Granger, in Jupyter Notebooks – a publishing format for reproducible computational workflows (2016).

80. M. Rocklin, in Proceedings of the 14th Python in Science Conference (2015), pp. 130–136.

81. K. Jordahl, J. V. D. Bossche, M. Fleischmann, J. Wasserman, J. McBride, J. Gerard, J. Tratner, M. Perry, A. G. Badaracco, C. Farmer, G. A. Hjelle, A. D. Snow, M. Cochran, S. Gillies, L. Culbertson, M. Bartos, N. Eubank, Maxalbert, A. Bilogur, S. Rey, C. Ren, D. Arribas-Bel, L. Wasser, L. J. Wolf, M. Journois, J. Wilson, A. Greenhall, C. Holdgraf, Filipe, F. Leblanc, geopandas/geopandas: v0.8.1 (Zenodo, 2020;https://zenodo.org/record/3946761).

82. A. Miles, J. Kirkham, M. Durant, J. Bourbeau, T. Onalan, J. Hamman, Zain Patel, Shikharsg, M. Rocklin, R. Dussin, V. Schut, E. S. D. Andrade, R. Abernathey, C. Noyes, Sbalmer, Pyup. Io Bot, T. Tran, S. Saalfeld, J. Swaney, J. Moore, J. Jevnik, J. Kelleher, J. Funke, G. Sakkis, C. Barnes, A. Banihirwe, zarr-developers/zarr-python: v2.4.0 (Zenodo,2020; https://zenodo.org/record/3773450).

83. B. Ripley, B. Venables, D. M. Bates, K. Hornik, A. Gebhardt, D. Firth, M. B. Ripley, Package ‘MASS.’CRAN Repos Httpcran R-Proj OrgwebpackagesMASSMASS Pdf (2013).

84. E. Paradis, S. Blomberg, B. Bolker, J. Brown, J. Claude, H. S. Cuong, R. Desper, Package‘ape.’Anal. Phylogenetics Evol. Version. 2(2019).

85. R. J. Hijmans, J. van Etten, raster: Geographic data analysis and modeling. R Package Version. 2, 15 (2014).

86. H. Stanke, A. O. Finley, A. S. Weed, B. F. Walters, G. M. Domke, rFIA: An R package for estimation of forest attributes with the US Forest Inventory and Analysis database. Environ. Model. Softw. 127, 104664 (2020).

87. E. P. Creeden, J. A. Hicke, P. C. Buotte, Climate, weather, and recent mountain pine beetle outbreaks in the western United States. For. Ecol. Manag. 312, 239–251 (2014).

88. S. Gillies, others, rasterio: geospatial raster I/O for {Python} programmers (2013;https://github.com/mapbox/rasterio).

